# Heterocellular coupling between amacrine cell and ganglion cells

**DOI:** 10.1101/328815

**Authors:** Robert E. Marc, Crystal Sigulinsky, Rebecca L. Pfeiffer, Daniel Emrich, James R. Anderson, Bryan W. Jones

**Affiliations:** Department of Ophthalmology & Visual Sciences, University of Utah, Salt Lake City, UT 84132; Signature Immunologics Inc., Torrey, UT 84775

## Abstract

All *superclasses* of retinal neurons display some form of electrical coupling including the key neurons of the inner plexiform layer: bipolar cells (BCs), amacrine or axonal cells (ACs) and ganglion cells (GCs). However, coupling varies extensively by *class*. For example, mammalian rod bipolar cells form no gap junctions at all, while all cone bipolar cells form class-specific coupling arrays, many of them homocellular *in-superclass* arrays. Ganglion cells are unique in that classes with coupling predominantly form heterocellular cross-class arrays of ganglion cell::amacrine cell (GC::AC) coupling in the mammalian retina. Ganglion cells are the least frequent superclass in the inner plexiform layer and GC::AC gap junctions are sparsely arrayed amidst massive cohorts of AC::AC, bipolar cell BC::BC, and AC::BC gap junctions. Many of these gap junctions and most ganglion cell gap junctions are *suboptical*, complicating analysis of specific ganglion cells. High resolution 2 nm TEM analysis of rabbit retinal connectome RC1 allows quantitative GC::AC coupling maps of identified ganglion cells. Ganglion cells classes apparently avoid direct cross-class homocellular coupling altogether even though they have opportunities via direct membrane touches, while transient OFF alpha ganglion cells and transient ON directionally selective (DS) ganglion cells are strongly coupled to distinct amacrine / axonal cell cohorts.

A key feature of coupled ganglion cells is intercellular metabolite flux. Most GC::AC coupling involves GABAergic cells (γ+ amacrine cells), which results in significant GABA flux into ganglion cells. Surveying GABA coupling signatures in the ganglion cell layer across species suggests that the majority of vertebrate retinas engage in GC::AC coupling.

Multi-hop synaptic queries of the entire RC1 connectome clearly profiles the coupled amacrine and axonal cells. Photic drive polarities and source bipolar cell class selec-tivities are tightly matched across coupled cells. OFF alpha ganglion cells are coupled to OFF γ+ amacrine cells and transient ON DS ganglion cells are coupled to ON γ+ amacrine cells including a large interstitial axonal cell (IAC). Synaptic tabulations show close matches between the classes of bipolar cells sampled by the coupled amacrine and ganglion cells. Further, both ON and OFF coupling ganglion networks show a common theme: synaptic asymmetry whereby the coupled γ+ neurons are also presynaptic to ganglion cell dendrites from different classes of ganglion cells *outside* the coupled set. In effect, these heterocellular coupling patterns enable an excited ganglion cell to directly inhibit nearby ganglion cells of different classes. Similarly, coupled γ+ amacrine cells engaged in feedback networks can leverage the additional gain of bipolar cell synapses in shaping the signaling of a spectrum of downstream targets based on their own selective coupling with ganglion cells.

## Introduction

Retinal ganglion cells are the signal outflow cells of the vertebrate retina: a network layer that integrates bipolar cell and amacrine cell signals and passes them to CNS targets. Like bipolar cells, most ganglion cells appear part of a unidirectional synaptic chain in not evidencing any direct feedback to the preceding input stage. However, early intracellular dye injection studies (Vaney, 1991;Vaney and Weiler, 2000;Vaney, 2002) showed intercellular dye diffusion patterns between ganglion and amacrine cells that have been interpreted as coupling mediated by gap junctions (e.g.Xin and Bloomfield, 1997b;Massey, 2008). Like synapses, gap junctions are extremely diverse structures serving intercellular signaling. The primary proteins of gap junctions are drawn from a large family of connexins with four transmembrane spanning domains and cytosolic domains that usually (not always) provide predominantly homotypic or bihomotypic binding even if the junctions are heteromeric (Li et al., 2008;Rash et al., 2013), and intracellular domains that mediate recognition and binding of other gap junction proteins. In general, it is thought that the peak open conductance of a single connexon is principally related to its pore diameter (this is not always true) with complex modulation enabled by a range of mechanisms (Ek-Vitorin and Burt, 2013;Hervé and Derangeon, 2013) including connexin phosphorylation (Pereda et al., 2013;O’Brien, 2017), methanesulfonate-analogue (taurine) binding (Locke et al., 2011), and many different adapter protein interactions (e.g. Zou et al., 2017).

Modes of coupling in the retina can be grouped into broad categories such as homocellular (coupling between the same “types” of cells) and heterocellular (coupling between different cell types). But what do we mean by “type” in the context of retina? Our terminology is based on computational classification theory where a *class* is the ultimate level of granularity (Marc and Jones, 2002). In this terminology, mammalian rod photoreceptors, blue cones, rod bipolar cells, A_II_ amacrine cells are all classes. In contrast, the categories of photoreceptors, bipolar, amacrine and ganglion cells are all *superclasses* as they contain collections of classes or larger intermediate groups often defined *ad hoc* (see Table S1). So what we really mean by heterocellular coupling is that it occurs between superclasses with clearly different morphologies. Homocellular coupling occurs within classes or a between intermediate groups with the same morphology. Thus CBb3n::CBb4 coupling, where:: denotes the presence of gap junctions between the pair, is homocellular (between bipolar cells) but is cross-class coupling (Table 1; also see Mills, 2001). As we will show, specific ganglion cells in the retina show common rules for heterocellular coupling with γ+ amacrine cells, ranging from none to extensive. And cells with strong GC::AC coupling so far show no proven in-class homocellular GC::GC coupling and certainly no cross-class coupling in the RC1 dataset of >1.4 million annotations. This makes them unique among retinal cells in largely avoiding homocellular in favor of heterocellular coupling.

**Table 1.**
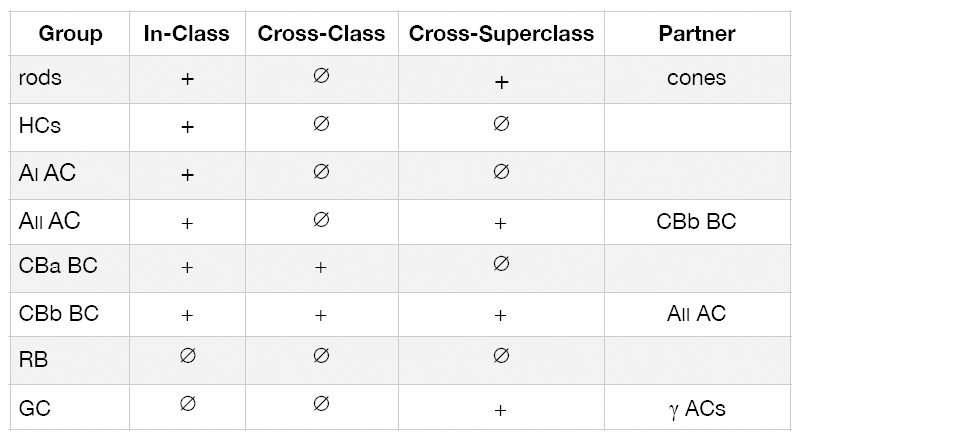
Patterns of retinal coupling: + exists, Ø null

|  | Homocellular |  | Heterocellular |  |
| --- | --- | --- | --- | --- |
| Group | In-Class | Cross-Class | Cross-Superclass | Partner |
| rods | + | ∅ | + | cones |
| HCS | + | ∅ | ∅ |  |
| A<sub>I</sub> AC | + | ∅ | ∅ |  |
| A<sub>II</sub> AC | + | ∅ | + | CBb BC |
| CBa BC | + | + | ∅ |  |
| CBb BC | + | + | + | A<sub>II</sub> AC |
| RB | ∅ | ∅ | ∅ |  |
| GC | ∅ | ∅ | + | γ ACs |

While we know quite a bit about the general patterns of GC::AC coupling from dye injection studies, the network frameworks for the specific partnerships involved and significance of coupling is elusive. There has been discussion about whether coupling leads maladaptive receptive field center expansion that would depress spatial resolution (Massey, 2008). However, two anatomical tools can assess the extent of coupling, enable precise definition of the partners and lead to more refined models of function: computational molecular phenotyping (CMP) and connectomics. While physiologic analyses will always be definitive arbiters of global network functionality, connectomics can resolve network topologies that physiology cannot (e.g. Lauritzen et al., 2016). CMP allows quantitative specification of the small molecule signatures of retinal neurons, especially ganglion cells (Marc et al., 1995). In mammals, many classes of ganglion cells are clearly coupled to GABAergic amacrine cells (γ+ amacrine cells), generating an intrinsic GABA signal superimposed on a classic high-glutamate, high-glutamine and low taurine ganglion cell signa-ture (Marc and Jones, 2002). As we will show, that signal is not unique to mammals. Using this signature and merging CMP with high-resolution connectomics allows us to trace the connectivity of specific identified retinal ganglion cells. To attack this problem directly we simply asked: what is the network embedding of GC::AC motifs? The broad conclusion for two specific ganglion cell classes is that heterocellular coupling enables diverse modes of network specificity depending on the topology of the coupled inhibitory network. The specific conclusion for the tON DS ganglion cell network is that excitation of the ganglion cell can lead directly to the inhibition of neighboring ganglion cells of differing classes through the tON DS GC::AC >_i_ GC chain. (where >_i_ denotes sign-inverting synaptic signaling)

## Methods

### Samples

Over 40 years our laboratory has collected retinal samples from over 50 vertebrate species spanning all classes. All euthanasia methods followed institutionally approved procedures, some of which changed over the years IACUC oversight evolved. Aquatic vertebrates were euthanized via cervical transection and double pithing (pre-1995) or sedated in 0.2% methanesulfonate prior to cervical transection (post-1995). Reptiles were similarly euthanized by cervical transection and double-pithing (pre-1995) or IP injection with 10% urethane followed by cervical transection. Mammals were euthanized by urethane overdose and thoracotomy (rabbits) or decapitation (pre-2014, mice), deep isoflurane anesthesia and thoracotomy or decapitation (2015); or Beuthanasia euthanasia and thoracotomy (rabbits, post-2015). The basic fixation method for all of them has been the same, as summarized in Marc et al. (1999b): 250 mM glutaraldehyde, 1320 mM formaldehyde in either cacodylate or phosphate 0.1M buffer pH 7.4, 3% sucrose, 1% MgSO_4_ or 1% CaCl_2_. All tissues were embedded in Eponate resins (Marc et al., 1978), serially sectioned at 100-250 nm onto array slides probed for small molecules (Marc et al., 1998), visualized by quantitative silver-immunogold detection (Marc and Jones, 2002), and imaged as described below. Some retinas were incubated for 10 minutes in either teleost saline (Marc et al., 1995) or Ames medium (Marc, 1999a,b) augmented with 5 mM 1-amino-4-guanidobutane (AGB) and either 1 mM NMDA or 0.05 mM kainic acid for excitation mapping of retinal ganglion cells.

### Immunocytochemistry

For the purposes of this paper, data from ≈20 years of postembedding immunocytochemistry were analyzed and summarized. The same protocols and antibodies were used for all analyses. It is important to note that post-embedding immunocytochemistry for glutaraldehyde-trapped amines or imines is idempotent: once the sample is fixed and embedded, no detectable changes in immunoreactivity occur, even over decades. The key marker for heterocellular coupling is 4-aminobutyrate (GABA) detected in post-embedding immunocytochemistry (Marc, 1999b) using YY100R IgG (RRID AB_2532061) from Signature Immunologics Inc. (Torrey, UT). Additional channels for cell classification (Marc et al., 1995; Anderson et al., 2009a;Anderson et al., 2011b) targeted AGB (B100R, RRID AB_2532053), glutamate (E100R, RRID AB_2532055), aspartate (D100R, RRID AB_2341093, glycine (G100R, RRID AB_2532057), glutamine (Q100R,RRID AB_2532059), and taurine (TT100R, RRID AB_2532060) from Signature Immunologics Inc. The activity tracer 1-amino-4-guanidobutane (AGB) is used to map both endogenous and exogenous ligand-driven glutamatergic signaling in single cells (Marc, 1999a;b;Marc and Jones, 2002;Marc et al., 2005;Anderson et al., 2011b). All IgGs were detected with silver-intensified 1.4 nm gold granules coupled to goat anti-rabbit IgGs (Nanoprobes, Yaphank, NY, SKU 2300), imaged (8-bit monochrome 1388 pixel × 1036 line frames) in large mosaic arrays with a 40x oil planapochromatic objective (NA 1.4) on a 100×100 Märzhäuser stage and Z-controllers with a QImaging Retiga camera, Objective Imaging OASIS controllers, and Surveyor scanning software (Anderson et al., 2011b; Lauritzen et al., 2016). Image quantification followed protocols described by Marc and Jones (2002), particularly histogram analysis which shows a distinct spectrum of signals for likely coupled retinal ganglion cells. Image analysis, histogram thresholding, object counts and spacing measures were performed using ImageJ 2.0.0-rc-43/1.51w (Rueden et al., 2017) in the FIJI Platform (Schindelin et al., 2012) and Photoshop CS6 (Lauritzen et al., 2016). Rendered neurons in RC1 were produced in Vikingplot (Anderson et al., 2011a;Anderson et al., 2011b) and VikingView (Lauritzen et al., 2016).

### Connectomics in rabbit retinal volume RC1

Connectome assembly and analysis of volume RC1 has been previously described (Anderson et al., 2009a;Anderson et al., 2011a;Anderson et al., 2011b;Lauritzen et al., 2012;Marc et al., 2013;Marc et al., 2014a;Lauritzen et al., 2016) and only key concepts expanded here. RC1 is an open-ac-cess rabbit retina volume imaged by transmission electron microscopy (TEM) at 2nm and includes 371 serial 70–90-nm-thick sections, capstoned and intercalated with optical sections containing small molecule signals (Anderson et al., 2011b). The retina was dissected from euthanized light-adapted female Dutch Belted rabbit (Oregon Rabbitry, OR) after 90 minutes (under 15% urethane anesthesia, IP) of photopic light square wave stimulation at 3Hz, 50% duty cycle, 100% contrast with a 3 yellow – 1 blue pulse sequence (Anderson et al., 2011b) with 13–16 mM intravitreal AGB. All protocols were in accord with Institutional Animal Care and Use protocols of the University of Utah, the ARVO Statement for the Use of Animals in Ophthalmic and Visual Research, and the Policies on the Use of Animals and Humans in Neuroscience Research of the Society for Neuroscience. Each retinal section was imaged as 1000-1100 tiles at 2.18 nm resolution in 16- and 8-bit versions, and as image pyramids of optimized tiles for web visualization with the Viking environment (Anderson et al., 2011a). Neural networks in RC1 have been densely annotated with the Viking viewer (Lauritzen et al., 2016), reaching over 1.4 million annotations of 3D rendered vol-umetric neurons, processes, pre- and postsynaptic areas, locations in the volume with subnanometer precision (Jensen and Anastassiou, 1995), and explored via graph visualization of connectivity and 3D renderings as described previously. The volume contains ≈ 1.5M annotations, 104 rod bipolar cells, > 190 validated cone bipolar cells, 300 amacrine cells and 20 ganglion cell somas. This density of annotations belies the additional work required to validate, classify and scale. Each annotation is a size and location entity coupled to a full metadata log (Anderson et al., 2011a) and has been validated by at least two tracing specialists; many have been revisiting 5-10 times, representing a total of 7 person-years of work. There is no automated tracing tool that doesn’t make more errors than a human; no tool (even our own Jagadeesh et al., 2013) that is so good that human cross-checking / validation / re-annotation is not necessary.

### Mining coupled ganglion cell networks

Candidate ganglion cell coupling networks in RC1 were visualized and annotated by identifying GABA+ ganglion cell somas and dendrites in Viking (connectomes.utah.edu, RRID:SCR_005986) in the intercalated GABA channels and by searching the RC1 database for coupling connections using network graph tools and database queries. All resources are publicly accessible via Viking and a range of graph and query tools are available at connectomes.utah.edu. All cells in this article are numerically indexed to their locations, network associations, and shapes. The data shown in every TEM figure can be accessed via Viking with a library of *.xml bookmarks available at marclab.org/GCACcoupling. Each cell index number in the RC1 database can be entered into different software tools for analysis, visualizations, or queries: Viking, Network Viz, Structure Viz, Info Viz, Motif Viz (Viz tools are based on the GraphViz API at graphviz.org, developed by AT&T Research, RRID SCR-002937), and VikingPlot developed by the Marclab; and VikingView developed by the University of Utah Scientific Computing and Imaging Institute. Further, Viking supports (1) network and cell morphology export into the graph visualization application Tulip (tulip.labri.fr) developed by the University of Bordeaux, France; (2) cell morphology for import into Blender (Blender.org, RRID SCR-008606); and (3) network queries for Microsoft SQL and Microsoft Excel with the Power Query add-in to use the Open Data Protocol (OData.org) to query connectivity features. More efficiently, we discover and classify coupling networks in Tulip with TulipPaths: a suite of regex (regular expression) based Python plug-ins for network queries (https://github.com/visdesignlab/TulipPaths, https://docs.python.org/2/library/re.html). Tulip networks can be directly exported from our connectome databases with a web query tool at connectomes.utah.edu and all data used in this article can be accessed via marclab.org/ <u>GCACcoupling</u> and the supplemental data file RC1_20180415_ALL-nw.tplx.

### Statistics

Small molecule signal comparisons across groups were done by both k-means clustering and histogram analysis using PCI Geomatica (Toronto, Canada) and Cell-Kit based on IDL (formerly ITT, now Harris Geospatial, Melbourne, FL) as described in Marc and Jones (2002) using. Various parametric and nonparametric analyses of feature sets (e.g. gap junction numbers, sizes) and power analyses were performed in Statplus:mac Version v6 (www.analystsoft.com/en/; RRID:SCR_014635) and R (www.r-project.org; RRID:SCR_001905).

## Results

### Phylogeny of ganglion cell::amacrine cell coupling

Diffusion of small molecule molecules like GABA through gap junctions readily identifies ganglion cells that may be coupled to GABAergic amacrine cells (Marc & Jones, 2002), and this GABA signature can be used to screen vertebrates for possible heterocellular GC::AC coupling. Specifically, cells in the ganglion cell layer with GABA signal histograms matching those of conventional amacrine cells (1-10 mM) are classified as displaced amacrine cells and those with intermediate signals (0.1-1 mM) are classified as provisionally coupled ganglion cells (see Marc and Jones 2002 for calibrations). In many species we are also able to correlate these intermediate GABA levels with classical high glutamate signals of ganglion cells and distinctly large ganglion cell sizes (e.g. Marc and Jones, 2002). Using the marclab.org tissue database we reviewed 53 vertebrate species spanning all vertebrate (Table S2) classes to assess the scope of potential coupling. Importantly, evidence of ganglion cell coupling occurs in *every vertebrate class*, even if other markers of comparative function vary: e.g. Müller cell GABA transport (limited to Cyclostomes, Chondrichthyes, Mammals and advanced fossorial ectotherms such as snakes), horizontal cell GABA transport (limited to most bony ectotherms) and horizontal cell GABA immunoreactivity (dominant in bony ectotherms and variable in mammals). The only vertebrate class we can say appears to clearly lack evidence of heterocellular GC::AC coupling is Testudines: turtles.

### Quantitative GABA immunocytochemistry

In every vertebrate class that shows coupling, the degrees of coupling and ganglion cell types involved are diverse. **Figure 1** shows the spectrum of GABA coupling signals in the rabbit ganglion cell layer just below the visual streak obtained by registering the glutamate (Fig. 1A) and GABA (Fig. 1B) channels of 2385 cells in the ganglion cell layer. The raw signals in Figure 1B reveal that GABA levels range from undetectable in many cells to levels that nearly match those of conventional amacrine cells, starburst amacrine cells in particular. But in between are a range of concentrations far lower than any GABAergic amacrine cell (Marc and Jones, 2002) but much higher than background. Our previous assessments of the selectively of the YY100R anti-GABA IgG (Marc and Jones, 2002) and competition assay results shown in Table S3 range from 10^4^-10^6^. Thus the intermediate values cannot be due to cross reactivity with any plausible alternate biomarkers (e.g. L-alanine, β-alanine, taurine, etc.) else they would have to present at levels of 1-100 M (100 µM low signal range × 10^4^-10^6^ cross-reactivity), which is physically impossible.

**Figure 1:**
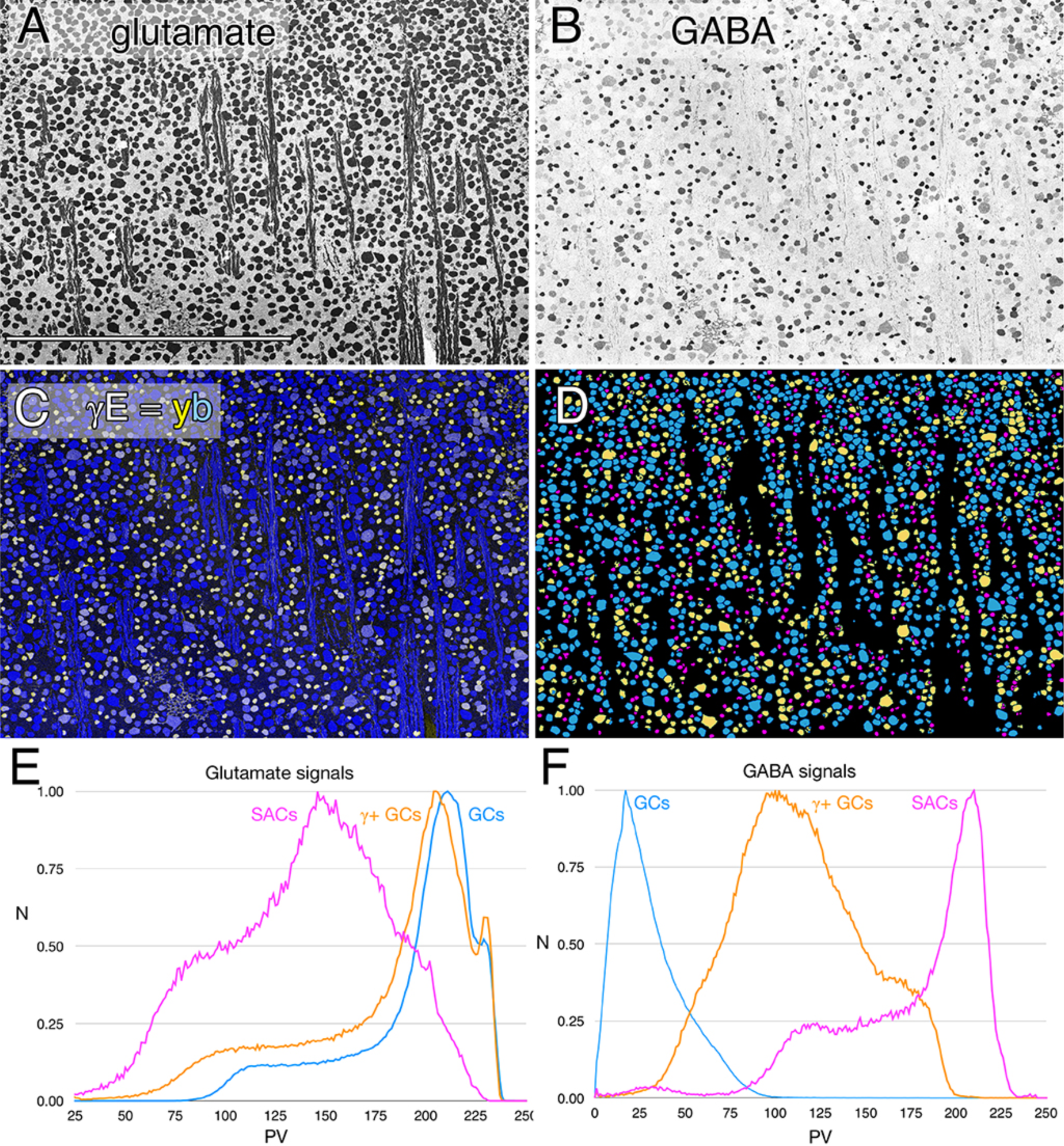
Glutamate and GABA colocalization in the rabbit ganglion cell layer; registered serial 200 nm horizontal sections through the plane of the ganglion cell layer with silver density visualization. A. Glutamate signals of ≈ 2100 cells, density mapped Scale, 0.5 mm.B. GABA signals of the same cells, density mapped. C. Intensity mapped registered channels with GABA signals encoded as a yellow channel (R + G) and glutamate signals as the blue channel. D. Theme mapped data produced through GABA histogram segmentation. Magenta: high GABA content (5-10 mM) population containing mostly starburst ACs and a few displaced ACs. Yellow: medium GABA content γ+ GCs (0.1-1 mM). Cyan: GCs with no measurable GABA content (<0.1 mM); small fragments represent portions of cross-sectioned cell dendrites. E. Glutamate histograms of peak normalized pixel number (N) vs pixel value PV for starburst ACs (SACs), GABA-positive GCs (γ+ GCs) and GABA-negative GCs (GCs). F. GABA histograms of peak normalized pixel number (N) vs pixel value PV for starburst ACs (SACs), GABA-positive GCs (γ+ GCs) and GABA-negative GCs (GCs). **Return**.

The intermediate ranges of GABA signals are associated with ganglion cell sizes ranging from some of the largest to some of the smallest ganglion cells (Fig. 1C) and the ganglion cell layer is separable by either clustering or histogram thresholding (Marc et al., 1995) into pure glutamate signal ganglion cells (uncoupled), γ+ ganglion cells (coupled) and starburst and minor displaced amacrine cell cohorts (Fig. 1D). Importantly, all γ+ ganglion cells show glutamate signatures indistinguishable from γ-pure glutamate ganglion cells (Fig. 1E), while starburst and other displaced amacrine cells display much lower glutamate contents similar to γ+ amacrine cell signatures in various vertebrate species (Marc et al., 1995;Marc, 1999a). The starburst amacrine cell cohort has a mean minimum feret diameter of 9.5 µm ± 1.8 µm (n=463), while the γ+ ganglion cell and ganglion cell cohorts are much more diverse since they represent multiple ganglion cell classes. More specifically, even with additional outlier cells (Marc and Jones, 2002) in the ON starburst amacrine cell cohort, the spacing is that of a unitary population with a mean nearest-neighbor spacing of 38.0 ± 14.5 µm and conformity ratio of 2.59 denoting ordered spacing with p < 10^−4^ (Cook, 1996). In contrast, the spacing data of γ+ and γ–ganglion cells are random, denoting mixed class aggregates. The diameters of γ+ ganglion cells range up to 35 µm in diameter, consistent with the sizes of OFF alpha ganglion cells (Marc and Jones, 2002) but existing within a more diverse cohort of medium sized cells overall (n=506, mean minimum feret diameter 15 ± 3.5 µm). Similarly, γ–ganglion cells range up to 36 µm in diameter consistent with the sizes of ON alpha ganglion cells, with a large cohort of medium sized cells (n=1049, mean minimum feret diameter 16.9 ± 3.6 µm). Basically, the amacrine cell cohort is unique in quantitative glutamate and GABA signatures, size and spacing while the γ+ ganglion cells and ganglion cells are not discriminable in glutamate signatures, size or spacing and are separable as an intermediate class only by the modest GABA signatures of γ+ ganglion cells. GABA in ganglion cells is basically a coupling signature.

Ganglion cell small molecule coupling is not unique to mammals. Teleost fishes represent the Actinopterygii, a vertebrate class with ≈ 400Mya divergence from class Sarcopterygii, while infraclass Teleostei is of even more modern origin (≈310Mya) with a massive post-Mesozoic, early Cenozoic expansion (Friedman, 2010) contemporaneous with mammalian speciation. The emergence of the cyprinids (goldfish and zebrafish) is extremely modern with estimated peak speciations in the Miocene and even early Holocene (Dubut et al., 2012), postdating the divergence of anthropoid primates (Pozzi et al., 2014). Arguably, comparing mammals and teleosts is one of the most diverse spans that could be conceived in assessing a putative synapomorphy such as coupling, with a last common ancestor in the Devonian. **Figure 2** shows GABA coupling patterns in the ganglion cell layer of the goldfish Carassius auratus. There are clearly different classes of ganglion cells with GABA coupling signatures that are below the amacrine cell signal range. As a control, the population of goldfish starburst amacrine cells forms a single signature cohort with GABA levels much higher than coupled ganglion cells. Their glutamate, GABA and kainate-activated AGB signals show that they form a distinct, monolithic, inseparable signature group that cannot be drawn from any other population, while presumably coupled and uncoupled ganglion cells have much weaker (or no) GABA signals and diverse signatures (Fig. 3). Furthermore, the starburst amacrine cell population shows a regular spacing pattern with a conformity ratio of 4.39, forming a statistical group distinct from a random distribution with p < 10-4 (Cook, 1996). This is a much higher precision mosaic than the rabbit, which is characteristic of teleost neuron patterning. The spacing of γ+ ganglion cells, in contrast, is completely random, suggesting that they represent a mixed population, as in rabbit. Similar results have been obtained in key model species such as mouse.

**Figure 2:**
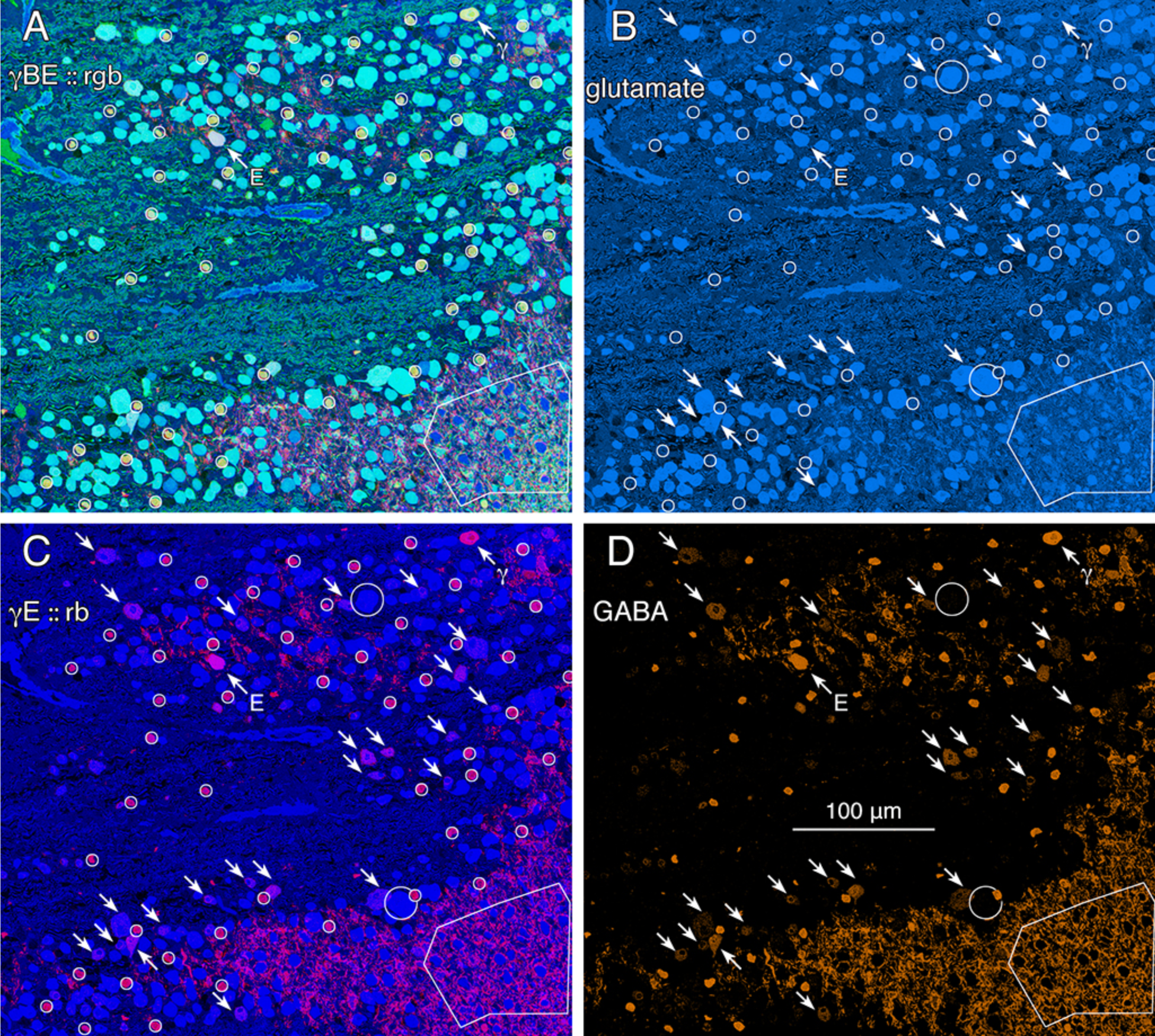
Glutamate and GABA colocalization in the goldfish ganglion cell layer; registered serial 200 nm sections with silver density visualization inverted to an intensity display (Marc et al., 1995). A. GABA (γ), AGB (B) and glutamate (E) signals visualized via γBE::rgb mapping. AGB permeation was activated *in vitro* with 50 µM kainic acid (KA) in the presence of 10 mM AGB in Hickman’s Teleost saline (Marc et al., 1995). This signature separates ON starburst ACs (circles) with an orange signal mixture (high GABA and AGB, repre-senting classic strong starburst AC KA responses) from cyan GCs (high glutamate and AGB, representing strong GC responses to KA), light blue GCs (high glutamate, low AGB, representing weak GC responses to KA), and deep blue spherical terminals of Mb ON bipolar cells (surrounded by a polygon), which lack ionotropic glutamate receptors. The E symbol indicates a high glutamate contents GC with modest γ+ and high AGB signals. The γ symbol indicates a low glutamate, high GABA non-starburst AC. B. Glutamate channel, intensity mapped as medium blue for visibility (R = 0, G ≈ 0.5B, B ≈ 0-240). Southeast arrows denote high glutamate GCs that also have significant GABA signals. C. GABA and glutamate channels mapped as γE ⟶ RB, revealing γ+ GCs as magenta cells. D. GABA channel mapped as orange for visibility (R ≈ 0-240, G ≈ 0.5R, B=0), clearly revealing weak GABA signals in a set of high glutamate GCs. Scale, 100 µm. **Return**.

**Figure 3:**
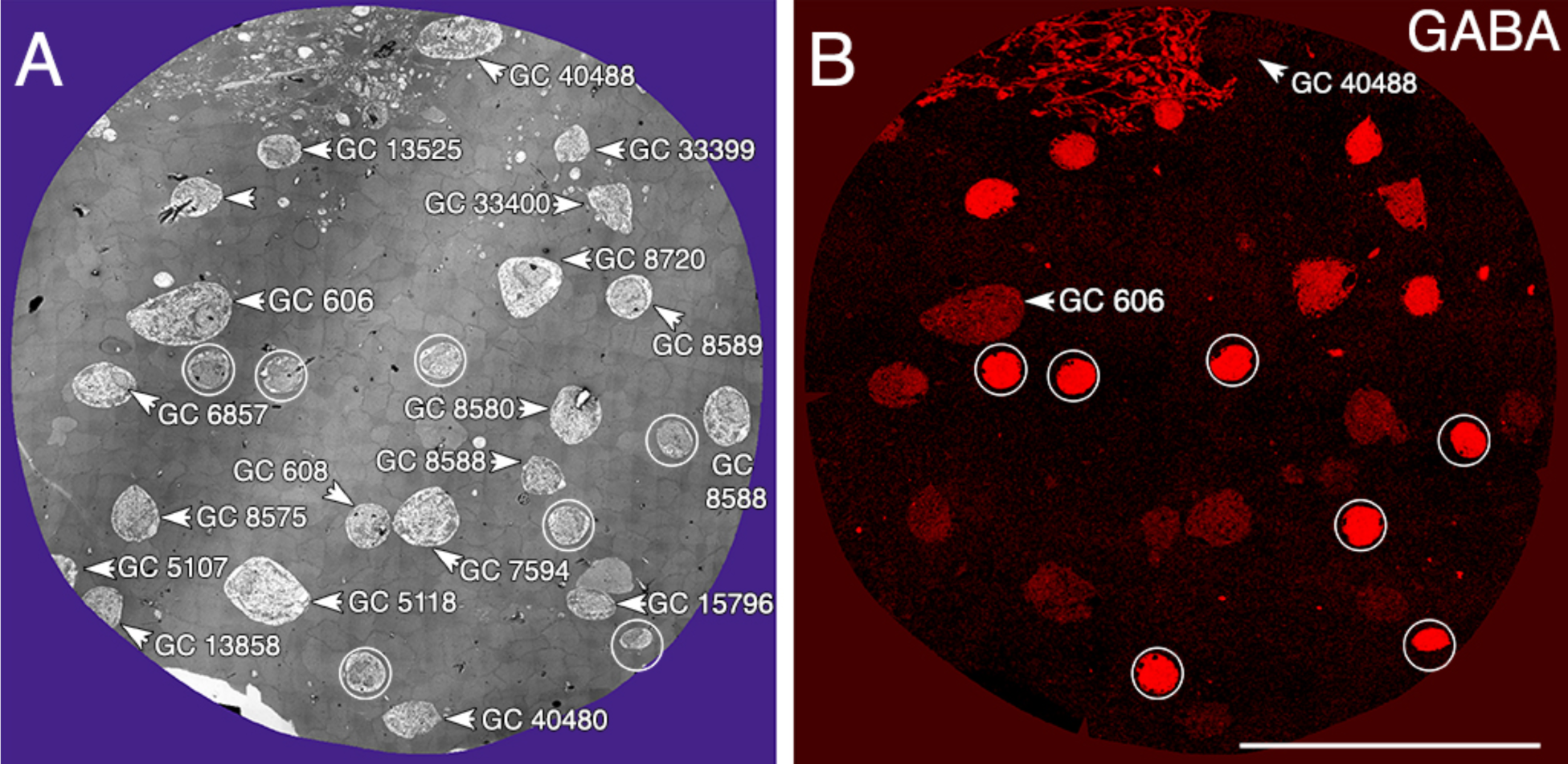
Ganglion Cell GABA colocalization in retinal connectome RC1. A. Slice 371 TEM image displaying somas of 20 GCs (numbered) and 7 ON starburst ACs (circled). GC 606 is the largest GC soma in the volume with major and minor diameters of 34 µm and 19 µm. B. Slice 371 TEM GABA channel (Anderson et al., 2011b) with γ+ GC 606 and γGC 40488 labeled. Scale, 100 µm. **Return**.

### Ultrastructural evidence

Tracer-coupling and GABA-coupling suggest widespread heterocellular GC::AC coupling in the vertebrate retina, and significant correlative evidence supports that view (Völgyi et al., 2013a). But what is the relationship between GABA-coupling and expression of gap junctions? This is where retinal connectomics can offer critical insights. The ganglion cell layer in rabbit retinal connectome RC1 contains the somas of 20 ganglion cells and 7 ON starburst amacrine cells (**Fig. 3A**). Several of the ganglion cells show significant levels of GABA (Fig. 3B), suggesting they may be coupled to γ+ amacrine cells. We have reconstructed the gap junction contact patterns of two major classes of γ+ ganglion cells. GC 606 is identifiable as a likely tON DS ganglion cell known to be dye-coupled to at least two classes of γ+ amacrine cells (Ackert et al., 2006;Ackert et al., 2009;Hoshi et al., 2011; Massey, personal communication), receives excitatory input exclusively from ON CBb bipolar cells and has a strongly GABA positive crescent-shaped soma in the ganglion cell layer (cf Ackert et al., 2006). The second cell class is a major dendrite from an OFF alpha ganglion cell 9787 that crosses the entire volume. Its soma lies outside the connectome since alpha ganglion cell dendritic arbors, often up to 0.9 mm in diameter, are much larger than the connectome volume diameter, 0.25 mm. However, the 1.5 µm diameter dendrite traverses the OFF layer, distinctly receives bipolar cell input exclusively from CBa bipolar cells and is also postsynaptic to every AII amacrine cell lobule it encounters (Marc et al., 2013; 2014) which is strongly diagnostic for OFF alpha cells. In addition, the dendrite traverses GABA-labeled slice z184 in the volume multiple times and is GABA positive. We traced most of the connections of both of these cells in RC1 and demonstrate that both are extensively coupled to unique sets of γ+ amacrine cells.

### GC 606

GC 606 is a large γ+ ganglion cell with a crescent shaped soma size of 35 µm maximum diameter, and it is strongly γ+ albeit at much lower concentrations than truly GABAergic amacrine cells such as ON starburst amacrine cells (Fig. 3). Its dendritic arbor spans the entire RC1 volume and is co-mingled with the arbors of γ+ interstitial axonal cells (IACs) and ON starburst amacrine cells (**Fig. 4**), and the excitatory synaptic input to GC 606 occurs from ON CBb bipolar cells that arborize distal (1-2 µm) to the ON starburst amacrine cell dendrites. The IAC is the γ+ PA1 polyaxonal cell ((Famiglietti, 1992;Wright and Vaney, 2004). GC 606 is indisputably an ON ganglion cell. There are no starburst amacrine cell inputs to GC 606 so it cannot be a classic sustained ON DS ganglion cell (Hoshi et al., 2009). As it is heavily coupled to γ+ amacrine cells it cannot be an ON alpha ganglion cell (Hu and Bloomfield, 2003). The size, arborization level, γ+ coupling and lack of starburst inputs are all consistent with the classification of GC 606 as a transient ON (tON) DS ganglion cell

**Figure 4:**
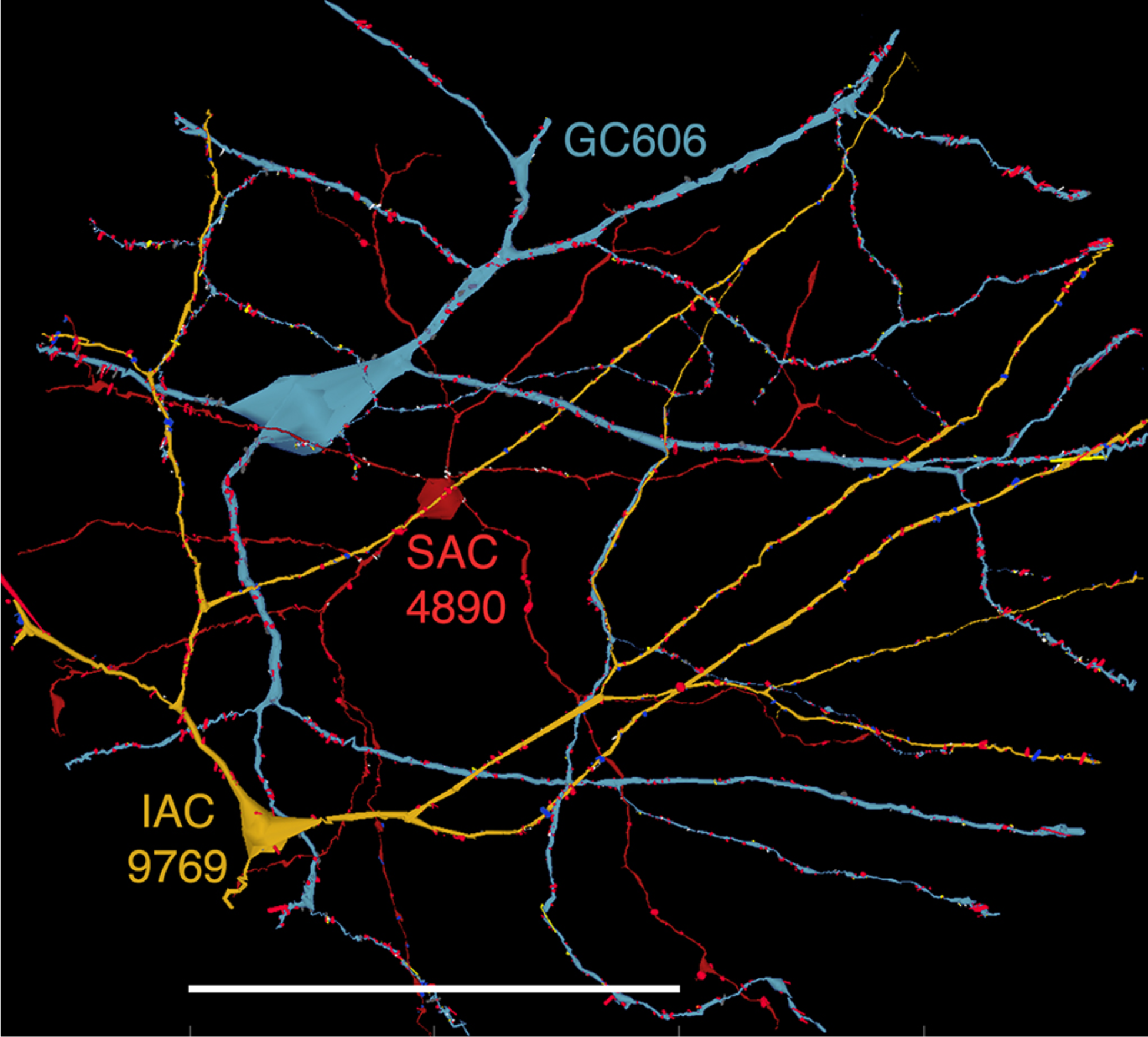
Commingled GC 606 (blue), starburst amacrine cell (SAC) 4890 (red) and IAC 9769 (gold) dendritic arbors in connectome RC1 visualized with VikingPlot rendering (Anderson et al., 2009b). Small dots are presynaptic specializations (blue), PSDs (red), gap junctions (gold), other non-synaptic features (grey, white). Scale, 100 µm. **Return**.

The initial stage of characterizing a neuron in a connectome is defining its excitatory, inhibitory and coupling drive (**Fig. 5**). The drive for GC 606 extracted by data queries from volume RC1 is summarized in Table 2 for 1267 validated contacts (supplemental file GC606-contacts.xlsx). As in previous analyses of the inner plexiform layer (Marc and Liu, 2000) synaptic drive is dominated by inhibition with ≈ 3 inhibitory synapses per excitatory input and 5.5 µm^2^ of inhibitory PSD area per µm^2^ of ribbon PSD. Despite the small size of ganglion cell gap junctions, they are abundant and the gap junction / ribbon ratio is 0.86 suggesting that nearly every ribbon input is near a coupling path. By measuring dendrite lengths of representative tON DS ganglion cells from Hoshi et al., (2011), we estimate that GC606 represents only 18% of the dendritic length of a complete ganglion cell. Thus each complete tON DS ganglion cell should receive ≈ 1440 ribbon synapses driving ≈ 35 µm^2^ of PSD area; ≈ 4350 inhibitory conventional synapses driving ≈ 290 µm^2^ of PSD area; and make ≈ 1270 gap junctions summing to 35 µm^2^ of copling area across the arbor. However this comprises only about 6% of the gap junction density in the IPL (Marc et al., 2014a) and since many of the gap junctions are suboptical, tracing them by fluorescence imaging (even super-resolution methods) could be challenging.

**Figure 5:**
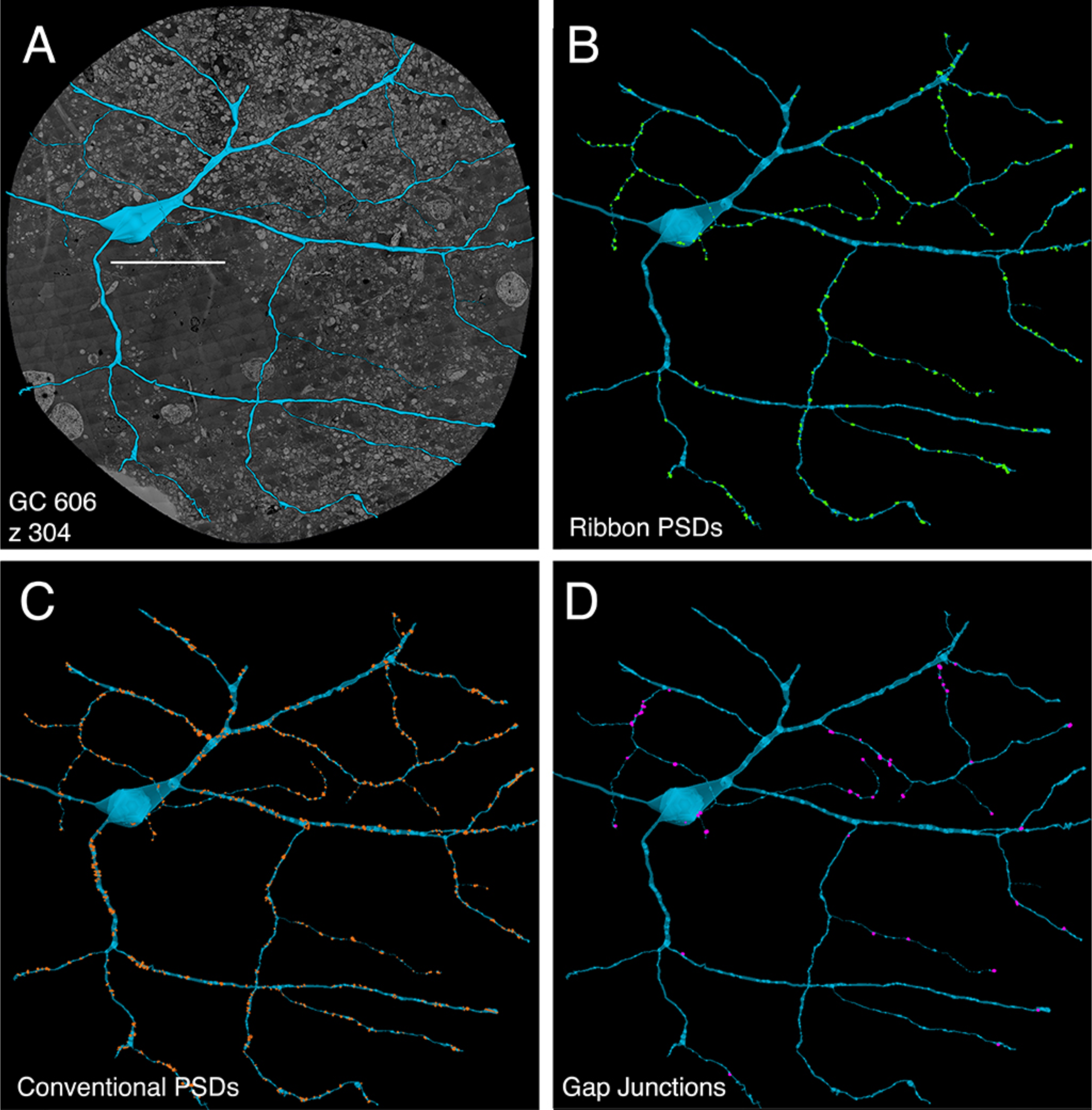
GC 606 superimposed on (A) connectome slice Z304 with separate displays of (B) excitatory ribbon synapse PSDSs, (C) inhibitory conventional synapse PSDs and (D) gap junctions each displayed at 2x their true diameters. Scale, 100 µm. **Return**.

**Table 2.**
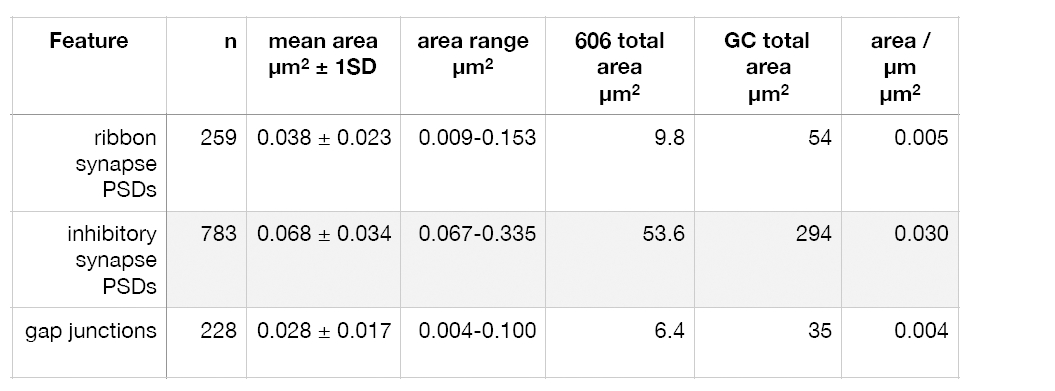
Contacts of GC 606

Excitation patterns are class-specific. GC 606 receives glutamatergic excitation exclusively from ON CBb bipolar cells as can be shown by querying the RC1 database with the TulipPaths plugin (see Methods): e.g. query “CBb.*, ribbon, GC ON” which returns all the ON CBb bipolar cell ribbon synapses onto specific ON ganglion cells from identified bipolar cells (**Fig. 6, 7**). Note that any single class of cone bipolar cells forms a closed-cell mesh of axonal field coupling (cf Mills and Massey, 1992). Of the 259 ribbon complexes that drive GC 606 in RC1, 54% are from one class of bipolar cells, CBb4w (Fig. 6A) and over 99% of the input *excludes* CBb5 bipolar cells which represent the primary drivers of ON starburst amacrine cells (**Fig. 7**). This is not due to stratification as both CBb4w and CBb5 bipolar cells have extensive overlap of their axonal arbors (Lauritzen et al., 2016). As a reference, IAC 9769 has extensive interaction with GC 606 and samples many of the same bipolar cells with an even narrower preference spectrum dominated by CBb4w and effectively excluding CBb5 (Fig. 7). In contrast, ON starburst amacrine cells contact a dif-ferent profile with over 90% of their inputs deriving from CBb5 and CBb6, and less than 1% from CBb4w (Fig. 7). While ON starburst amacrine cells make numerous synapses onto ganglion cell dendrites in the RC1 volume, they make no synapses onto either GC 606 or IAC 9769, nor do they receive any synapses from IAC 9769.

**Figure 6:**
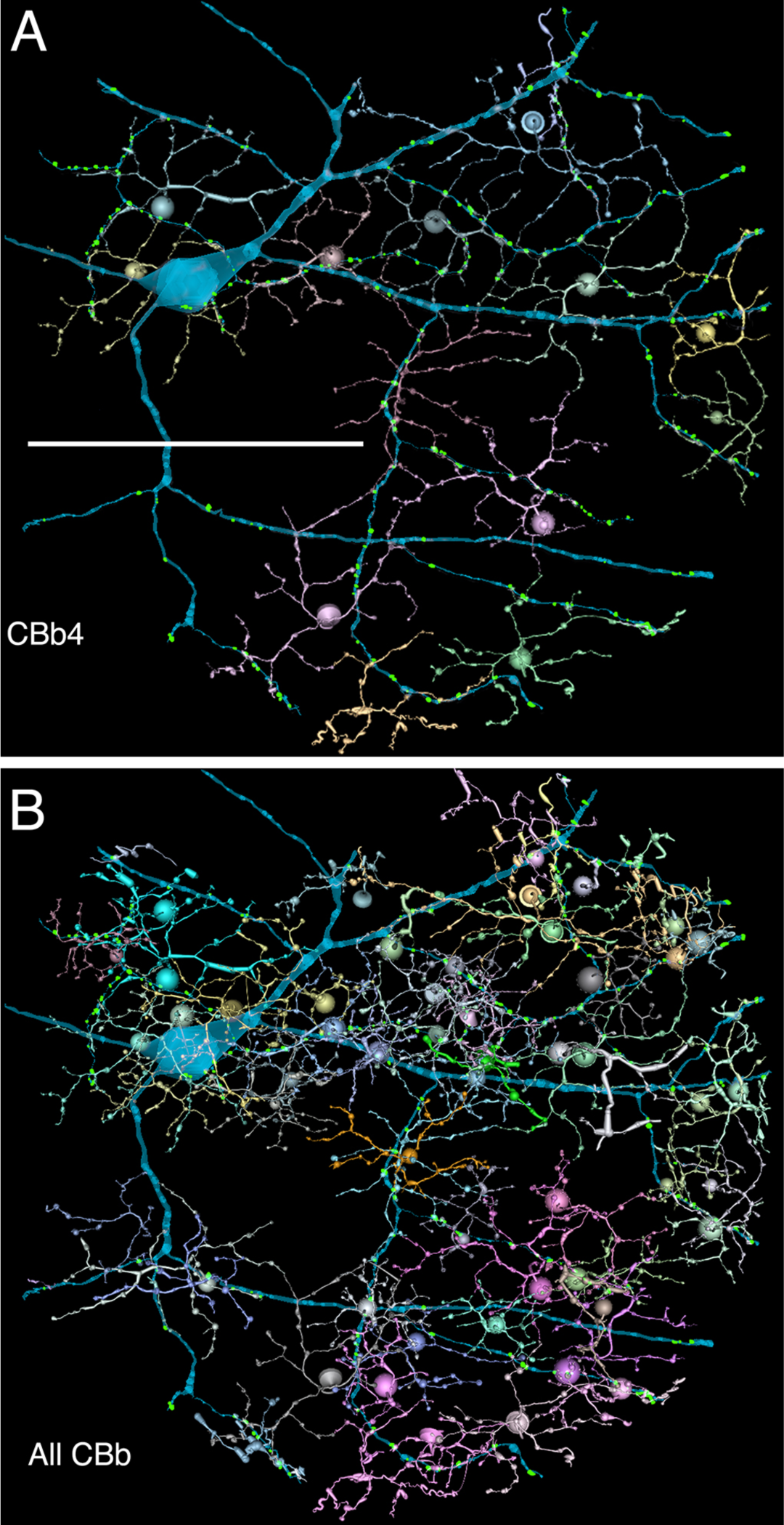
GC 606 and its bipolar cell input. A. The dominant synaptic ribbon drive (58%) arises from a single class, CBb4w, a coupled homocellular network of cone bipolar cells. Each cell is colored individually. B. The entire CBb input cohort to GC 606. Scale, 100 µm. Combined VikingPlot and VikingView rendering (Lauritzen et al., 2016). **Return**.

**Figure 7:**
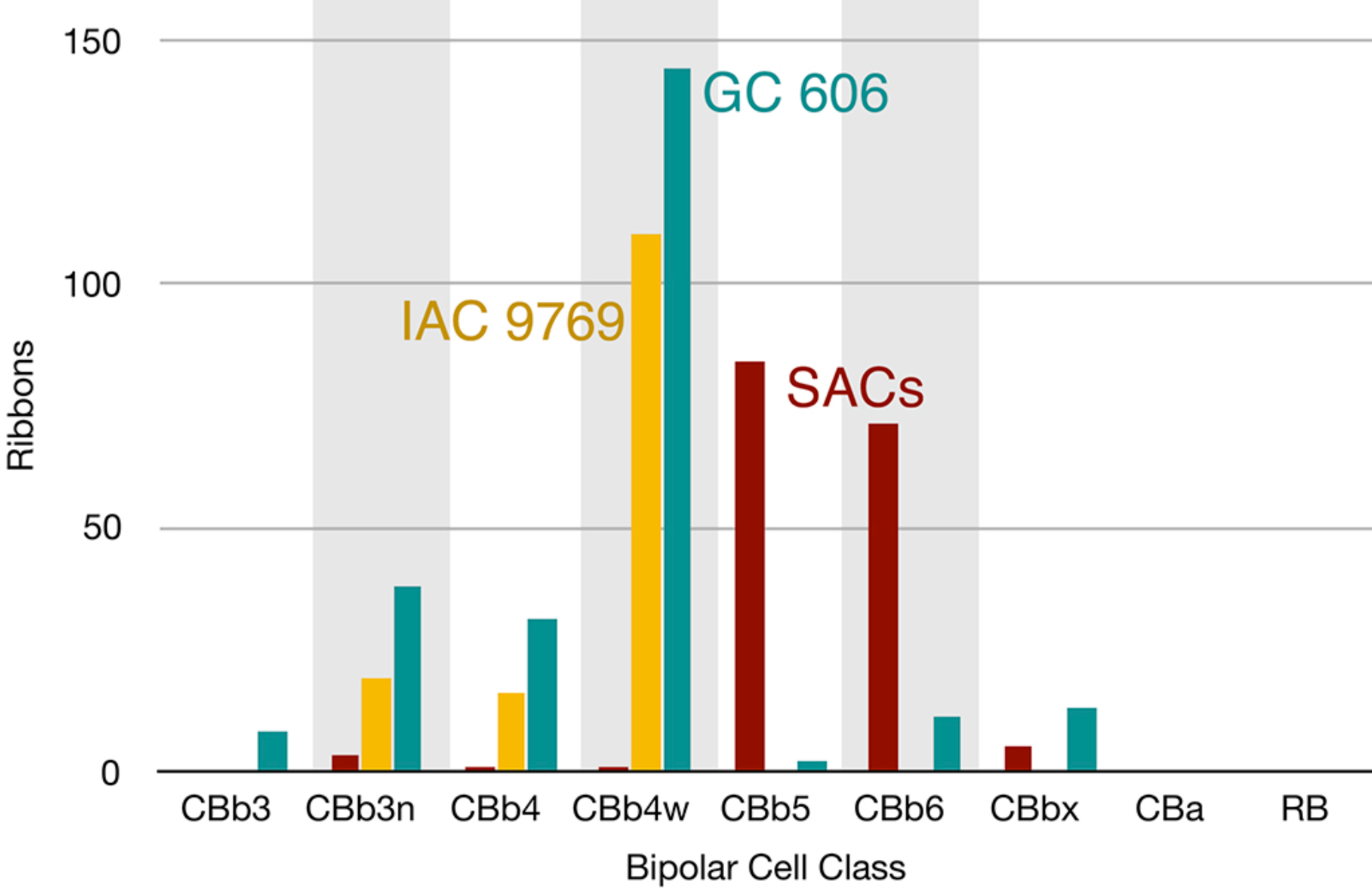
The classes of bipolar cell input to GC 606 (Cyan, n = 247), IAC 9769 (gold, n =145) and all ON SAC dendrites (red, n = 165) in RC1. Ordinate: number of synaptic ribbons from each class. Abscissa. All bipolar cell groups, including ON cone bipolar cell classes CBb3, CBb3n, CBb4, CBb4w, CBb5, CBb6, the aggregate OFF cone bipolar cell superclass (CBa) and the rod BC class. CBbx cells are ON cone bipolar cells from the volume margins with insufficient reconstruction to allow identification. **Return**.

In addition to its extensive CBb bipolar cell input, GC 606 also collects 783 conventional synapses from amacrine cells. Of those that are neurochemically identified, 33 have been mapped to definitive γ+ amacrine cells and only 2 to glycinergic amacrine cells as they traverse GABA or glycine reference slices (see Anderson et al., 2011b).

The key feature that distinguishes GC 606 is its extensive and obvious coupling with amacrine cells and IACs (**Fig. 8**). In parsing the entire ganglion cell arbor, it is clear that it makes abundant small gap junctions with amacrine cell-like processes (Table 2). Overall, the entire GC 606 arbor in RC1 makes 228 gap junctions of which 61 have been successfully traced to specific source amacrine cells or IACs. The mean gap junction diameters for the entire cohort (181 ± 56 nm) are not significantly different from those of the identified amacrine cell subset (paired homoscedastic T-test, p = 0.43, dof = 284). The diameter range is 72 - 357 nm, and many gap junctions are thus sub-optical. The morphology of retinal gap junctions is characteristic of vertebrate CNS, yielding multilaminar profiles at 0.27 nm/pixel resolution with spacing identical to those reported by Marc et al.(1988) using ≈ 0.1 nm resolution on film. All but two of the GC gap junctions in the entire volume RC1 are associated with amacrine cell processes and every GC::AC gap junction that is associated with a complete soma or traverses a reference slice arises from a GABAergic amacrine cell. **Figure 9** illustrates the arbor of GC 606 and its overlap with IAC 9769 (Fig. 9A) and a set of γ+ amacrine cells (Fig. 9B). Key locations where representative gap junctions are formed are marked as A1, A2, B1, B2, etc. and displayed in **Figure 10**.

**Figure 8:**
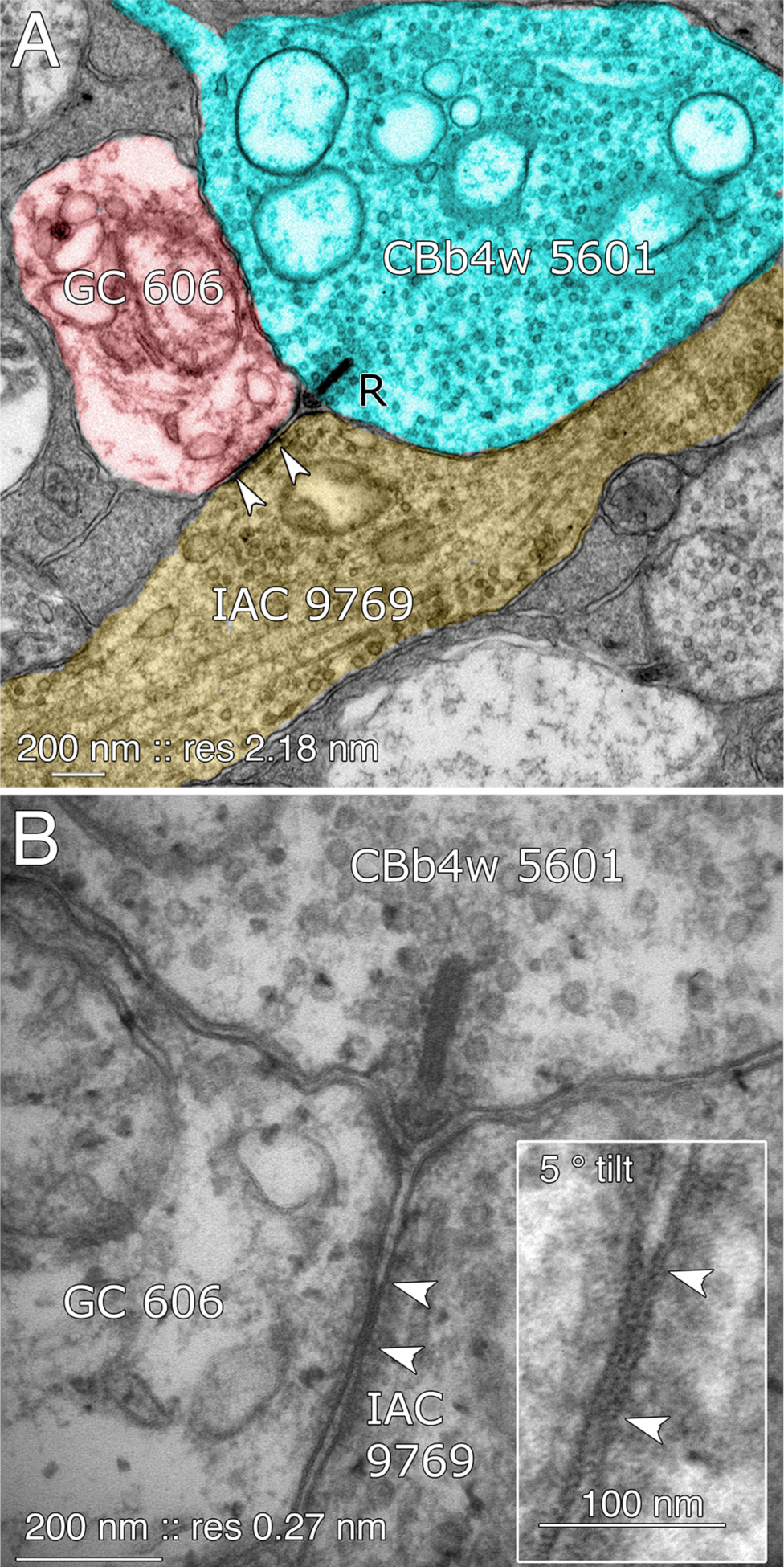
Coupling between GC 606 and IAC 9769. A. Connectome RC1 image of CBb4w 5601 (cyan) providing dyadic synaptic ribbon (R) input to GC 606 (red) and IAC 9769 (yellow). A large gap junction between GC 606 IAC IAC 9760 is visible as a unique dense line over the apposed membranes of the two cells (bracketed by arrowheads). This is the basic identification schema for identifying gap junctions in the volume at its native 2.18 nm/pixel. Note that the gap junction can be “zoomed” to subpixel image levels in practice for an-notating it (Anderson et al., 2011a). B. TEM reimaging of the same gap junction and ribbon complex visualized at high resolution (0.27 nm/pixel) and goniometrically tilted 5° to optimize the multilaminar gap junction structure (inset). Reprinted by permission from (Marc et al., 2013). **Return**.

**Figure 9:**
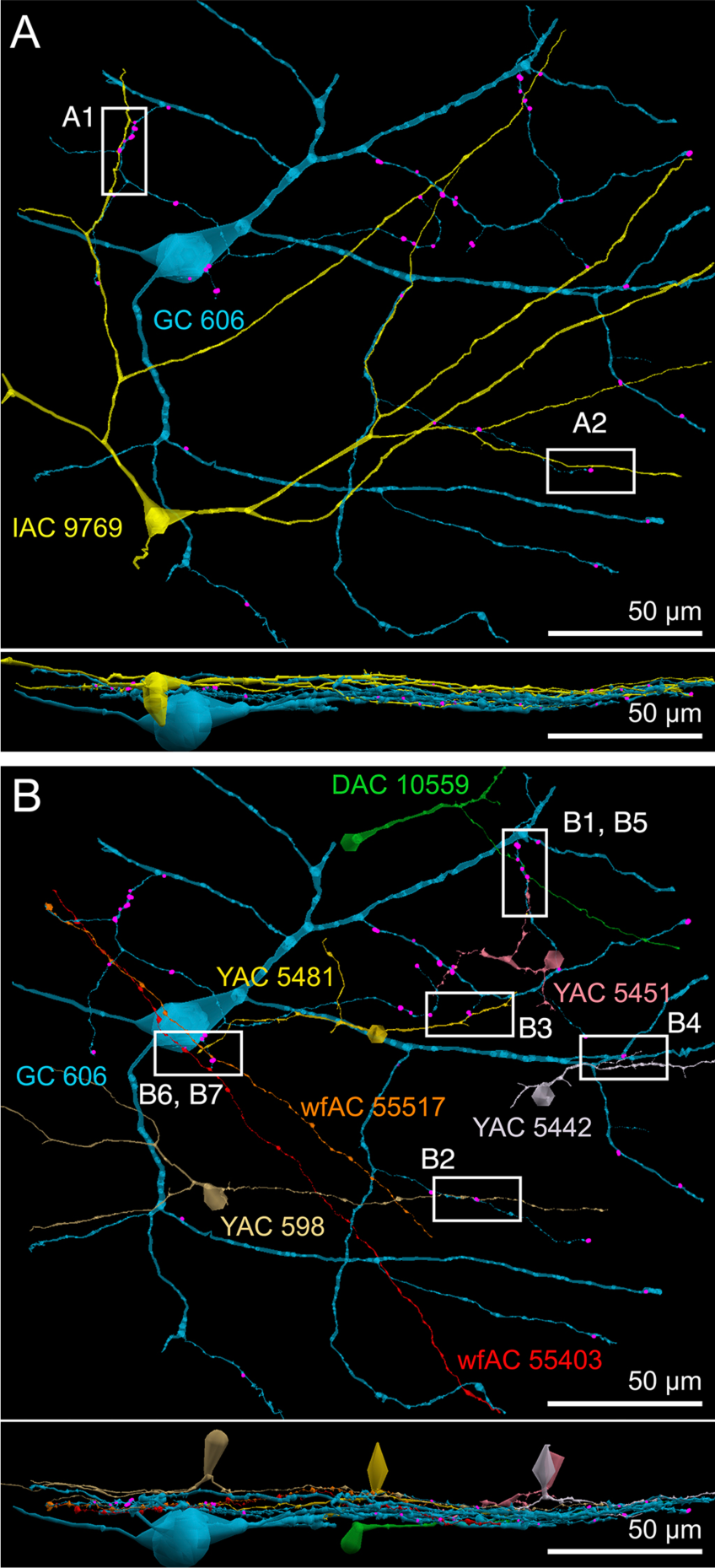
Selected sites of coupling between inhibitory neurons and GC 606. A. Two loci of coupling (A1, A2) between IAC 9769 and GC 606 viewed as a horizontal field. Lower image, vertical overlay. B. Seven loci of coupling between displace amacrine cell (DAC) 10559; γ+ amacrine cells with somas in the RC1 volume (YACs) 5481, 5442, 5481 and 598; and wide-field γ+ amacrine cell (wfAC) processes 55403 and 55517 arising from somas outside the volume. Horizontal and vertical overlays. High resolution analyses of these loci are shown in Figure 10. **Return**.

**Figure 10:**
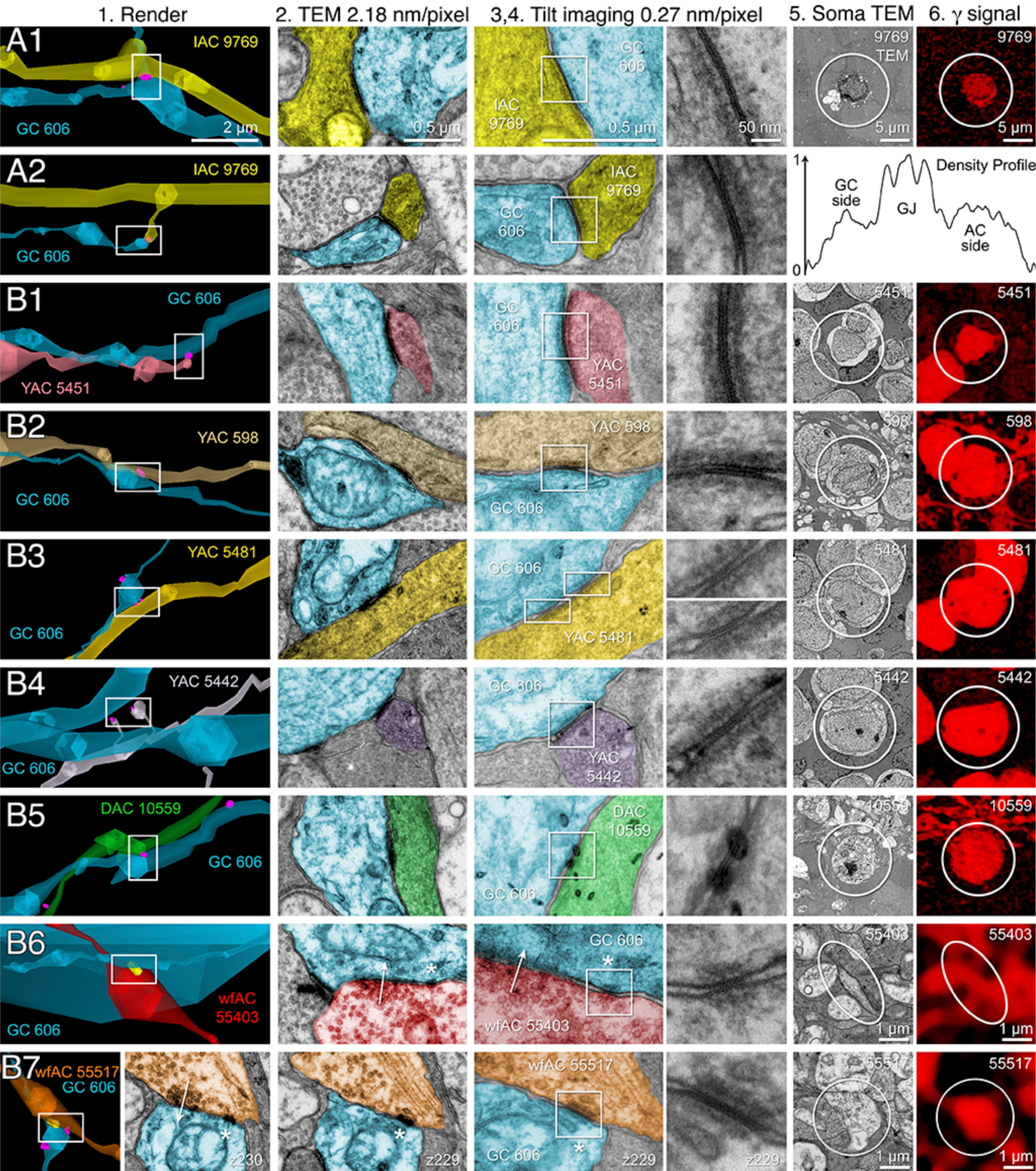
High-resolution analysis of coupling loci in Figure 9 imaged as: Column 1, VikingPlot renders; Column 2 TC1 native TEM at 2.18 nm/pixel; Columns 3 and 4, Goniometric reimaging at 0.27 nm/pixel; Column 5, soma or major process TEM; Column 6, GABA (γ) signal from the nearest intercalated CMP channel (Anderson et al., 2011b). A1. Gap junction between GC 606 (cyan) and IAC 9769 (yellow). A2. Gap junction between GC 606 (cyan) and IAC 9769 (yellow). Inset in columns 5,6 show a normalized plot of membrane density spanning the entire junction starting from the paramembranal density in GC 606, crossing the trilaminar zone and ending in IAC 9769 (ImageJ). B1. Gap junction between GC 606 (cyan) and γ amacrine cell YAC 5451 (pink). B2. Gap junction between GC 606 (cyan) and γ amacrine cell YAC 598 (tan). B3. Gap junction between GC 606 (cyan) and γ amacrine cell YAC 5481 (yellow). B4. Gap junction between GC 606 (cyan) and γ amacrine cell YAC 5542 (lavendar). B5. Gap junction between GC 606 (cyan) and displaced γ amacrine cell DAC 5451 (green). Note that the lamination can be visualized through the inadvertant stain debris in column 4. B6. Gap junction between GC 606 (cyan) and wf γ amacrine cell wfAC 55403 (red). B7. Gap junction between GC 606 (cyan) and wf γ amacrine cell wfAC 55517 (orange). Return.

While the arbor of IAC 9769 coarsely intertwines with GC 606 at several loci, *fasciculation doesn’t correlate with the occurrence of gap junctions*, which typically appear at brief crossing points where the processes align for less than a few µm and even then do not occur along the apparent alignment (Fig. 9A). From a TEM perspective, gap junctions occur at loci where gaps in suboptical glial processes expose the target, similar to axonal ribbons in bipolar cells (Lauritzen et al., 2012). IACs are not the only γ+ neurons that couple with GC 606. A set of conventional amacrine cells driven by CBb bipolar cells are also coupled to GC 606 (Fig. 9B). While their reconstructed fields are too limited to classify them all, they mostly appear to be wide-field (wf) γ+ amacrine cells, and there may be two or three classes that couple to GC 606. Representative validated gap junctions from IAC 9769 and the other γ+ amacrine cells are shown in Fig. 10. At high resolution, it is clear than most gap junctions are not fasciculations but rather crossings (Fig. 10 Column 1). The resolution of RC1 (2.18 nm/pixel) is sufficient to reliably detect gap junctions and measure their areas (Fig. 10 Column 2) but is not adequate for complete validation as occasional adherens junctions can mimic gap junctions in oblique view (Marc et al., 2014b). High resolution (0.27 nm/pixel) reimaging with goniometric tilt allows validation of gap junctions by visualizing their characteristic multilaminar density profiles (Fig. 10 Columns 3,4 and inset).Finally, all of these coupled neurons are GABAergic (Fig. 10 Columns 5,6). Of course it is not possible to reimage every structure in every grid, but of the more than 2000 partneridentified gap junctions tagged in RC1 at 2.18 nm/pixel resolution, ≈ 20 have proven to be mistaken adherens junctions (<1% error).

The real advantage of TEM connectomics database analysis is that we can take additional network hops and ask what the roles of the coupled interneurons might be. Every cell that is coupled to GC 606 is exported as a *.tlp format and its embedding network displayed in the Tulip framework (tulip.labri.fr). All the amacrine cells coupled to GC 606 show that their excitatory drive arises from CBb3, CBb3n, CBb4 and/or CBb4w bipolar cells and that they avoid the CBb5 and CBb6 cells that drive starburst amacrine cells and sustained ON and transient ON-OFF DS ganglion cells. While many ON γ+ amacrine cells are predominantly feedback amacrine cells that target CBb bipolar cells, this specific cohort also shows direct feedforward to ganglion cell classes other than GC 606. In general, the density of feedback synapses in the ON cone bipolar cell network in the entire volume RC1 appears ≈ 3:1 higher than feedforward synapses: 2359 feedback bipolar cell, 336 feedforward ganglion cell and 564 feedforward amacrine cell synapses. This lumped analysis masks the exceptional specificity of various well-known cells. For example, the cohort of rod bipolar cell-driven A_I_ amacrine cells in RC1 make 837 feedback synapses into bipolar cells and 0 feedforward synapses to either ganglion cells or amacrine cells. This makes their network error rate lower than 0.12%. In contrast, IAC 9769 has a strongly reversed bias at >10:1 feedforward:feedback, targeting 38 amacrine cells, 13 ganglion cells and only 3 cone bipolar cells (**Fig. 11**). Feedforward does not load the upstream network like feedback does, allowing for strong channel isolation even if the interneuron is involved in feedback. An excellent example is ON γ+ AC 598 (Fig. 9B) which engages in both feedforward and feedback, transferring sign-conserving coupled signals from GC 606 via sign-inverting GABA synapses to another ON GC (**Fig. 12**).

**Figure 11:**
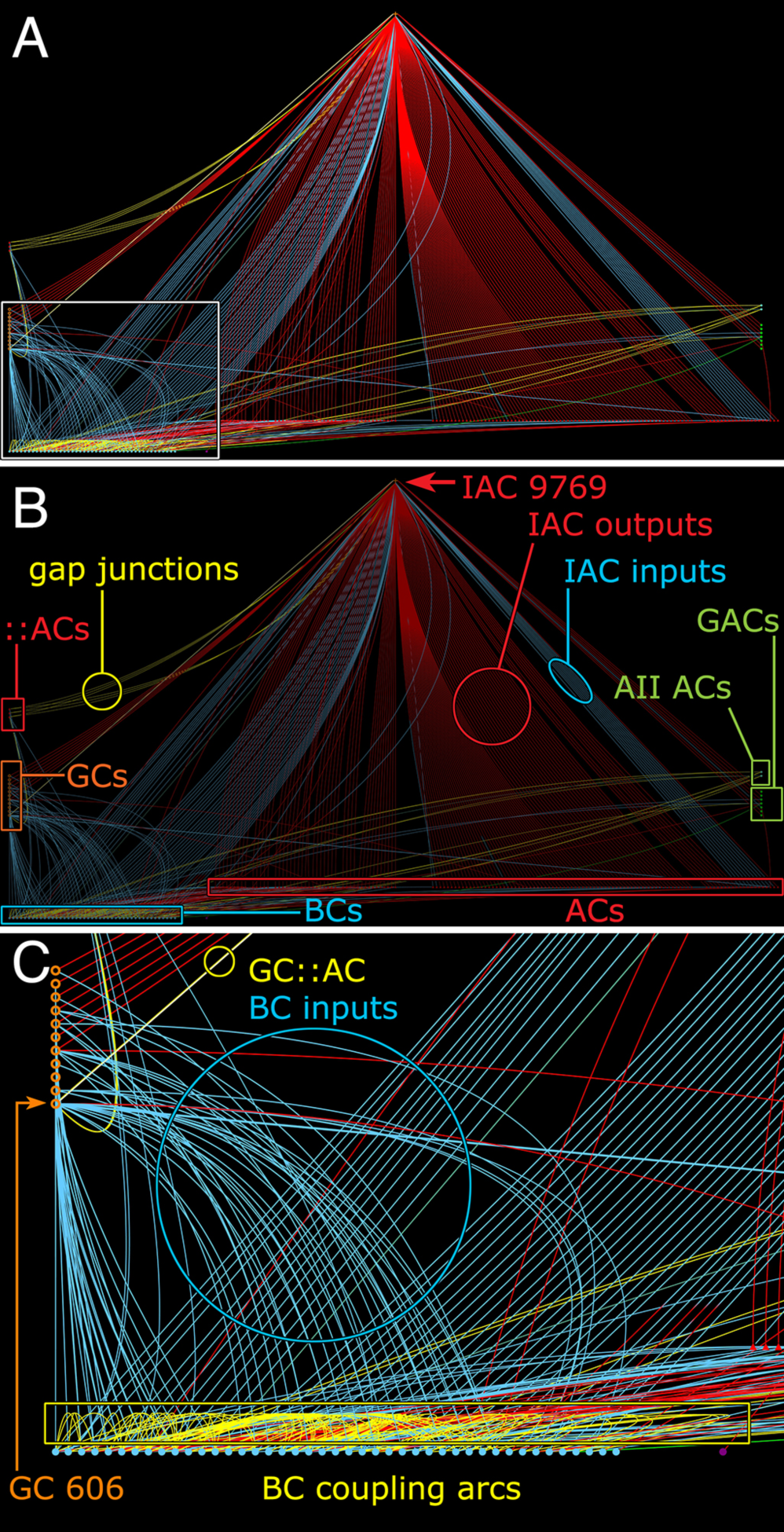
Connectivity of IAC 9769 to amacrine, bipolar and ganglion cells. Tulip query bundled (i.e. multiple synaptic paths between loci are bundled as single lines) dendrogram.A. Annotation-free view with panel C inset marked (box). B. Key: Red symbols, IAC, γ+ ACs and unidentified ACs; green symbols, A_II_ and GACs (glycinergic amacrine cells; blue symbols, bipolar cells; orange symbols, ganglion cells; small red symbols at right, IAC coupled::ACs (coupled amacrine cells); red lines and arcs, synaptic outflow from IAC 9769; blue lines and arcs, synaptic input to IAC 9769 and instances of bipolar cell; yellow lines and arcs, gap junctions. C. Enlargement of inset in A. GC 606 is strongly coupled to IAC 9769 (circled yellow edge, GC::AC) and other γ+ amacrine cells (yellow arc). Massive coupling networks exist among ON cone bipolar cells (yellow box). **Return**.

**Figure 12:**
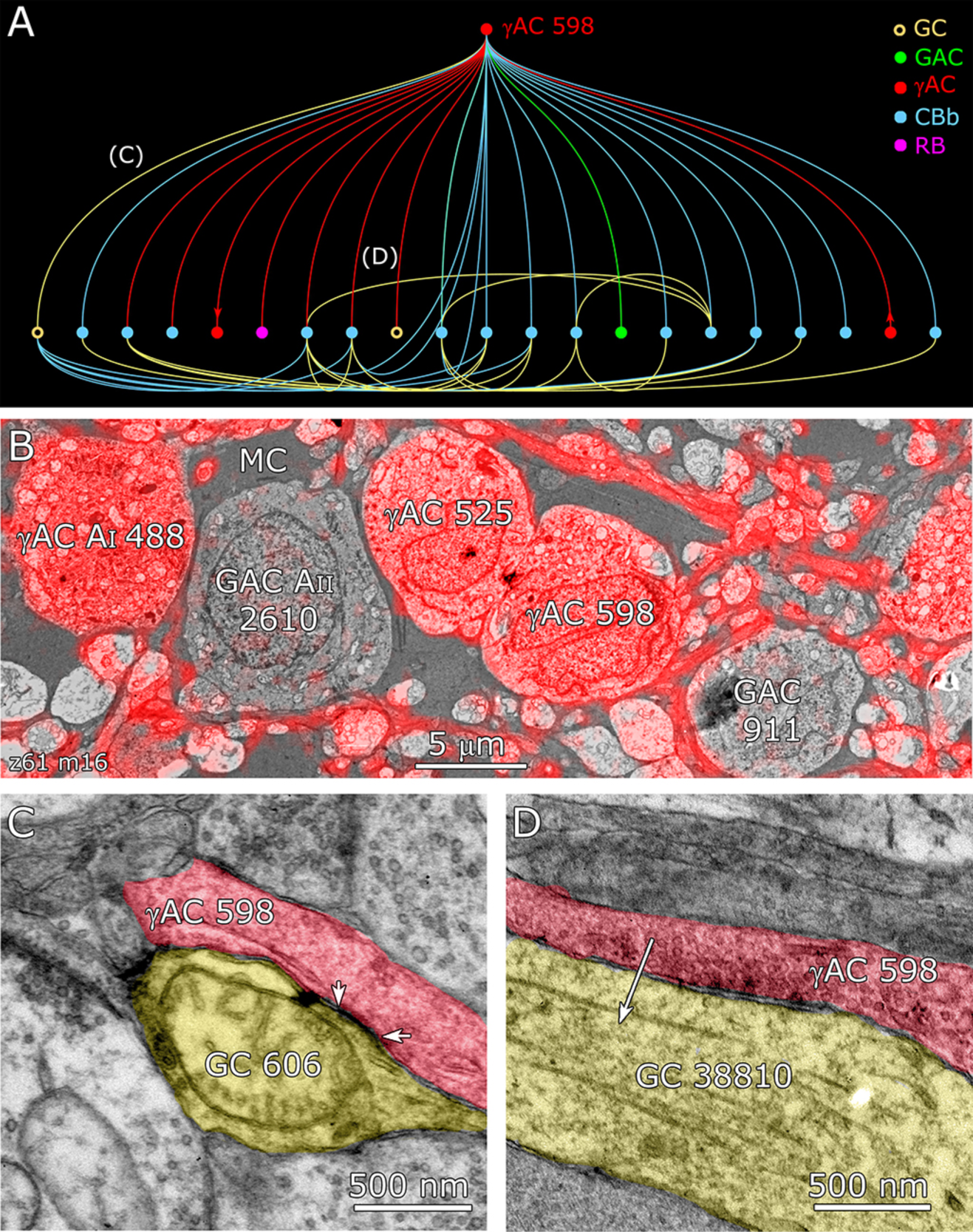
Coupling flow from GC 606 through γ amacrine cell 598 to multiple targets. A. Tulip query bundled dendrogram plot of all the sources and targets of amacrine cell 598: GC yellow circles, ganglion cells: GAC green dots, glycinergic amacrine cells; red dots γACs, GABAerigc amacrine cells; cyan dots CBb, ON cone bipolar cells; magenta dot RB, rod bipolar cell. Line color denotes the presynaptic source. Arrows denote presynaptic source in γAC to γAC paths. Each line represents a bundle of synaptic lines. B. Validation of GABAergic identity for AC 598. C. A gap junction between AC 598 (red) and GC 606 (yellow) delimited by arrowheads. D. Synapse from AC 598 (red) to GC 38810 (arrow). **Return**.

The coupled set of γ+ IACs / ACs and additional unclassified ACs form over 200 gap junctions with GC 606 *in the RC1 volume* implying that the complete cell forms over 1000 gap junctions, comprising a massive coupling path between the inhibitory and excitatory networks of the retina. Cross-class inhibitory feedforward driven by coupling to GC 606 converges on pure ON GCs (ID 606, 7594) and ON-OFF GCs (ID 5107, 6857, 15796). GC 7597 is also γ+ coupled (Fig. 3) albeit at lower levels than GC 606 and none of the GC 606-coupled amacrine cells appear to couple with GC 7597. ON-OFF GC 5107 is uncoupled and γ-while GC 15796 is very weakly γ+ and GC 6857 is strongly coupled. Thus feedforward inhibition does not appear to discriminate classes. We can summarize this chain as: GC1::AC >_i_ GC2 (where class 1 ≠ class 2, i.e. they are *disjoint* sets).

Other ganglion cells receiving feedforward are too incomplete to classify as they arise from outside the volume and it is impossible to connect branches to exclude mixed polarity inputs. Those with pure OFF inputs remain a possibility (e.g. ID 28950). If we use the rough scaling for size obtained in Table 2, a target ganglion cell could receive at least 200 inhibitory synapses via a single amacrine or axonal cell, driven by a coupled ganglion cell of a different class. This must be a vast underestimate since we cannot trace the majority of the coupled processes that arise from outside the volume. Finally, we have found no proven homocellular gap junctions between ganglion cells consistent with findings in mouse retina (Völgyi et al., 2013b). There are 2 candidate junctions out of many thousands of gap junctions in the volume, but we cannot validate them as ganglion cells. That does not mean they do not exist in rabbit since the RC1 volume is too small to ensure discovery of coupling between the small overlap zones of ganglion cells of the same class. However, the massive scope of heterocellular coupling suggests that even if GC::GC coupling occurs in homocellular overlap zones, it must be quantitatively much less significant than GC::AC coupling, realistically on the order of 0.4% based on our current encounter rates.

### GC 9787

Among the full cohort of ganglion cells, OFF alpha ganglion cells in the rabbit retina are characterized by a number of key features. First, in peripheral retina (rabbit volume RC1) they are among the largest of retinal ganglion cells with very large, simple dendrites of 1-2 µm diameter, dendritic arbors of ≈ 500-900 µm, somas approaching 30 µm in diameter and extensive heterocellular coupling to amacrine cells (Xin and Bloomfield, 1997a;Marc and Jones, 2002;Peichl et al., 2004). The somas can protrude deeply into the inner plexiform layer and insert large dendrites into the OFF layer of the inner plexiform layer. Additionally, they receive extensive input from both OFF (CBa) cone bipolar cells and A_II_ ACs (Kolb and Famiglietti, 1975;Marc et al., 2014a). Due to the sparse but uniform coverage of OFF alpha ganglion cells, we presumed that the largest crossing dendrite of the OFF layer in RC1 was the most probable candidate: GC 9787 (**Fig. 13**). GC 9787 and its input CBa bipolar cells stratify in the proximal half of the OFF stratum, while reference ON-OFF ganglion cells e.g. GC 5107 have their dendrites and input OFF bipolar cell terminals in the most distal portion of the IPL. Because the single unbranched dendrite is so large and represents a cell with a much larger field than nearby bistratified diving GC (Lauritzen et al., 2012) and even tON DS ganglion cells, this clearly excludes classical X-type sustained GCs and a range of W-type cells, even those that are γ+ coupled. Beyond size, four features suggest that the single large dendrite crossing the volume arises from an OFF alpha ganglion cell. First, the dendrite has a γ+ signature, as expected from heavily coupled OFF alpha ganglion cells (**Fig. 14**). Second, it collects inputs only from a subset of OFF bipolar cells (mostly CBa2), especially at multi-ribbon, large PSD sites (**Fig. 15**), avoiding CBa1 and CBa1w bipolar cells and axonal ribbons of ON bipolar cells in the OFF layer. In contrast dendrites of bistratified diving ganglion cells traverse the OFF layer, avoid CBa bipolar cells and selectively collect OFF-layer axonal ribbons from ON CBb bipolar cells (Lauritzen et al., 2012). GC 9787 also appears to form large PSDs (up to 600 nm diameter) only at pre-synaptic ribbon sites (Fig. 15) while completely avoiding conventional (ribbonless) bipolar cell synapses which are, in fact, quite common in the OFF layer and formed by the same bipolar cell instances onto different targets (Anderson et al., 2011b;Marc et al., 2013). For example, bistratified ON-OFF ganglion cell 8575 collects two *conventional ribbonless* OFF bipolar cell synapses for every OFF ribbon it contacts.Third, GC 9787 receives conventional synapses from every AII AC it encounters, six cells in all across the volume (**Fig.16**). Finally, it forms distinct gap junctions with amacrine cells (**Fig. 17**), including monostratified OFF YACs such as AC 7859 (Fig. 14A). While we cannot verify that every coupled process is γ+, there is no evidence that any glycinergic amacrine cell in RC1 is coupled to either GC 9787 or GC 606 in RC1. While we previously identified a single class of glycine and GABA coupled GC in the rabbit retina (Marc and Jones, 2002), we have not yet encountered a valid instance of glycinergic amacrine cell coupling to ganglion cells in RC1.

**Figure 13:**
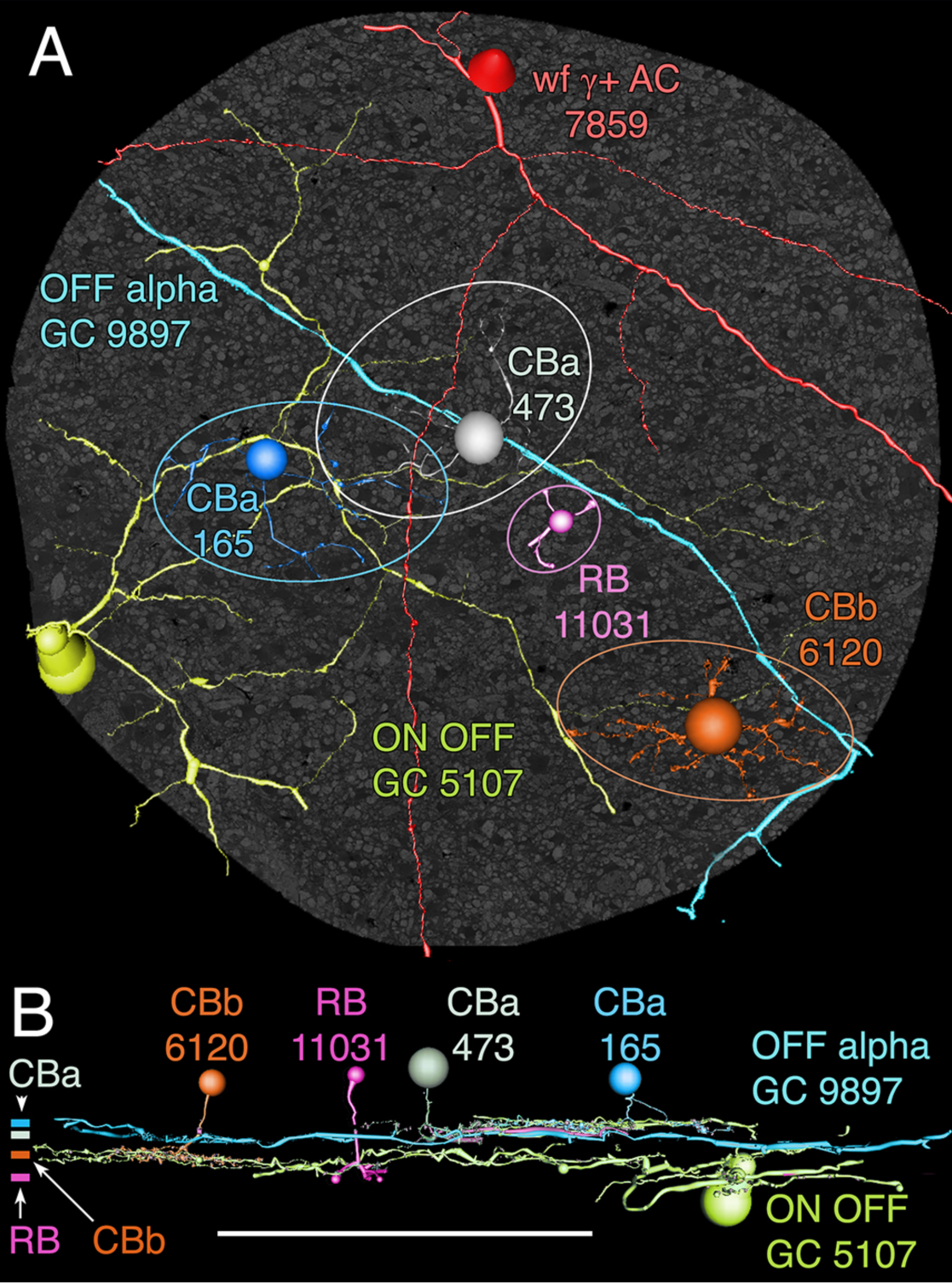
OFF alpha ganglion cell candidate dendrite GC 9897 crosses the connectome volume. A. Horizontal view of OFF alpha GC 9897 (cyan) dendrite in comparison to ONOFF GC 5107 arbor (yellow-green) and four reference bipolar cells. ON-OFF GC 5107 is driven by both ON cone bipolar cells (e.g. CBb5 6120 tangerine) and OFF cone bipolar cells (CBa 165 blue) that bracket the cone ON-OFF layers and are distal to the rod bipolar cell layer (e.g. RB 11031, magenta). GC 9897 is driven by a separate set of more proximal OFF cone bipolar cells (e.g. CBa 473 grey). Ellipses delimit bipolar cell axonal fields. B. Vertical view displaying the separate strata for rod (RB), ON cone (CBb) and OFF cone (CBa) bipolar cells. CBa 473 that drives OFF alpha GC 9897 is proximal to the OFF CBa 165 and similar bipolar cells that drive GC 5107. **Return**.

**Figure 14:**
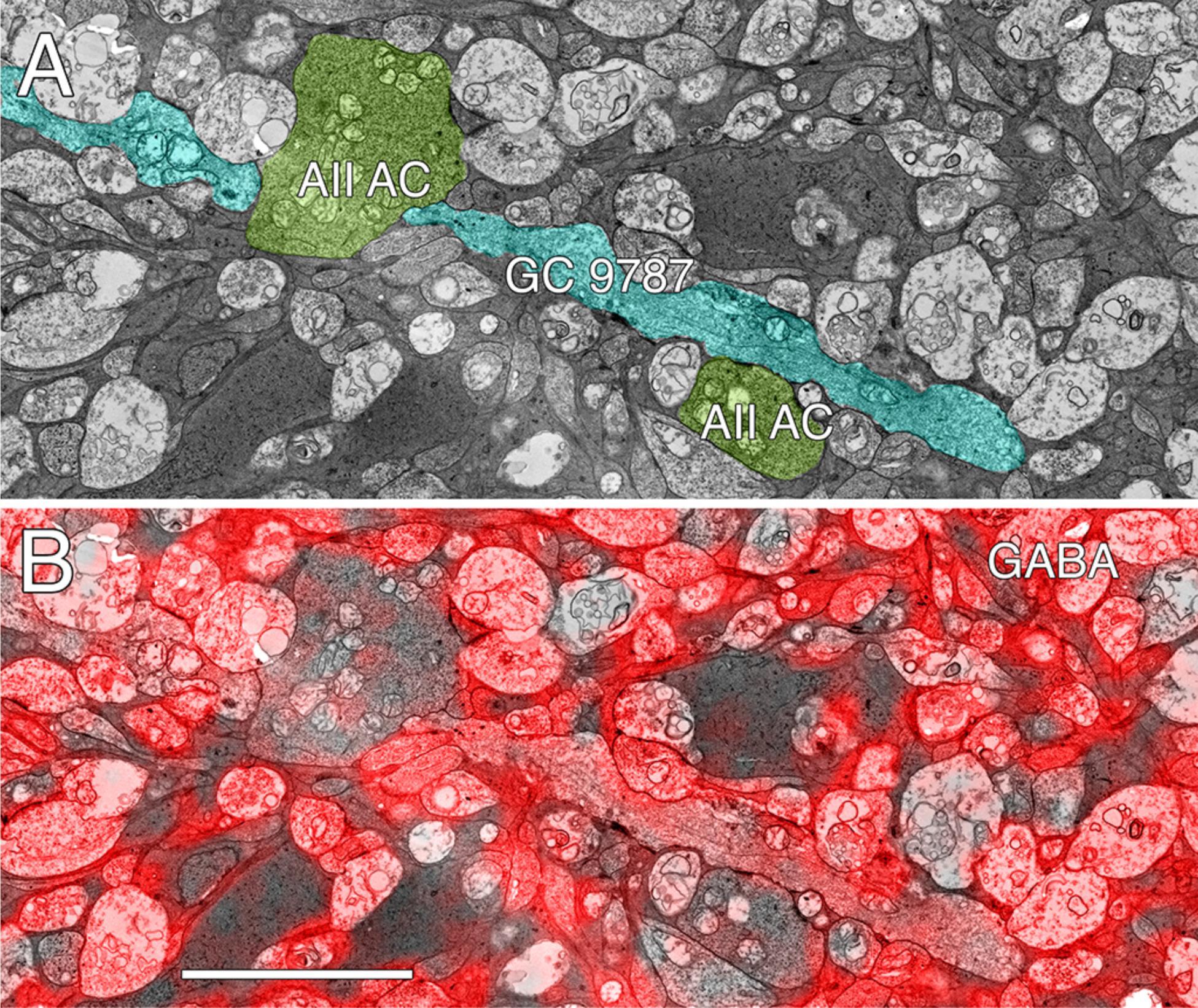
GABA-coupled signals in GC 9787. A. Single TEM slice z184 containing a segment of GC 9787 (cyan) flanked by A-II AC lobules (yellow-green). B. The same slice z184 TEM image with the neighboring intercalated GABA overlay showing positive colocalization. Scale 5 µm. **Return**.

**Figure 15:**
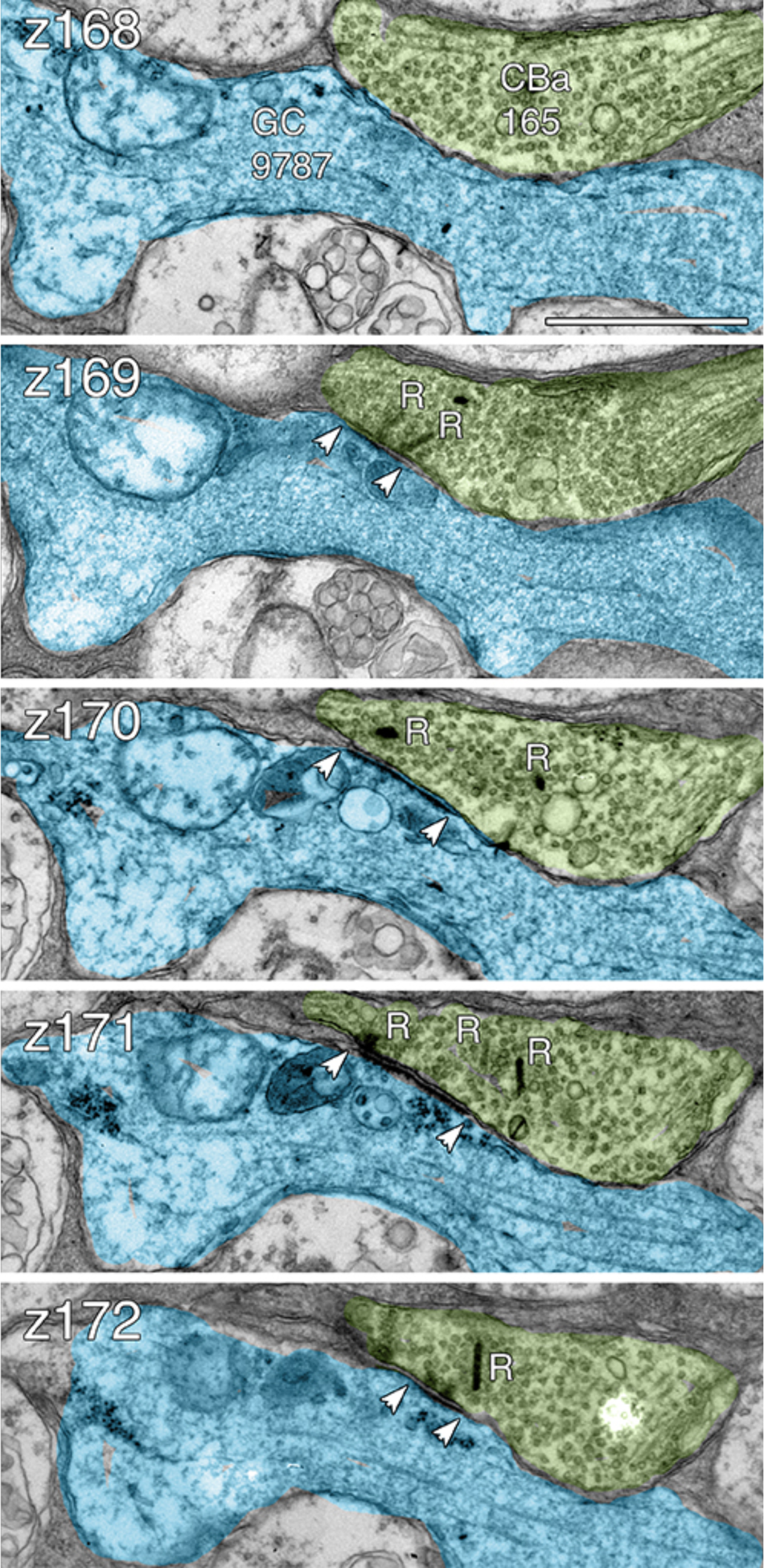
Multi-ribbon OFF cone bipolar cell inputs to GC 9787. Five serial TEM sections (z168-z172) are coded for GC 9787 (cyan) and CBa 165 (green). Over a span of 280-300 nm, at least 4 bipolar cell synaptic ribbons (R) dock presynaptically across from a single large postsynaptic density (bracketed by arrowheads). Arrows denote direction of synaptic flow (pre ⟶ post). Scale 1000 nm. **Return**.

**Figure 17:**
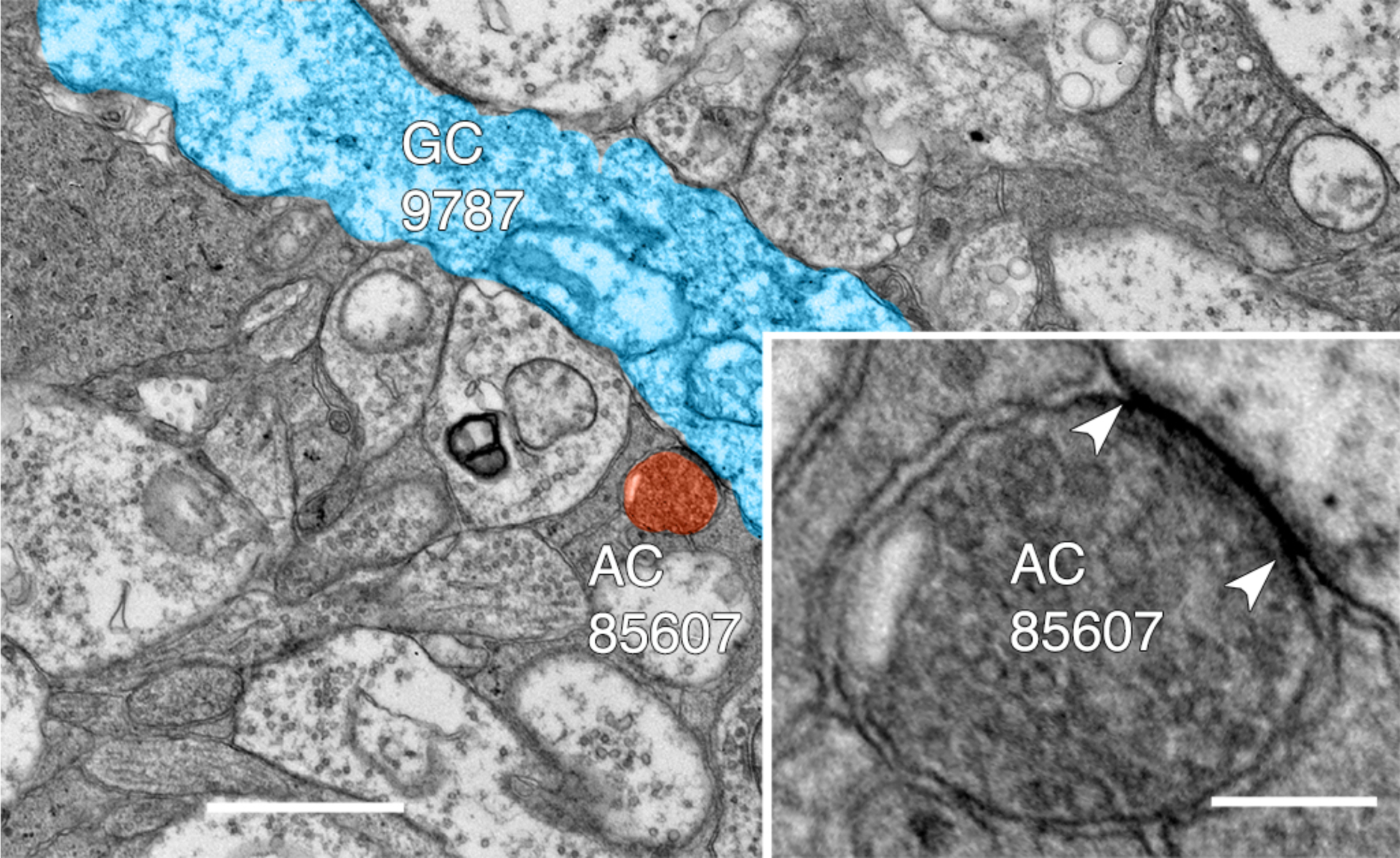
Typical gap junction between GC 9787 (blue) and an amacrine cell (AC 85607) at a crossing, non-fasciculated junction. Scale 1000 nm. Inset. High resolution image of the gap junction, showing characteristic gap occlusion. Scale 100 nm. **Return**.

Except for the southeast margin of the volume, GC 9787 is a smooth unbranched dendrite very unlike the topology of GC 606 and represents only ≈ 0.3 mm of length. The entire passage of GC 9787 through the OFF layer collects 13 gap junctions with an area of 9142 nm^2^ / µm of dendrite length, which is ≈ 46% of the gap junction density of GC 606. The moments and range of size of gap junctions formed by GC 9787 (≈ 37000 ± 19000 nm^2^) trend larger, but the sample is not significantly different from those formed by GC 606 (≈ 28000 ± 18000 nm^2^) assessed by either parametric (unpaired, heteroscedastic t-test; Ftest) or non-parametric (Komologorov-Smirnov measures). As power is set by the smallest sample group, however, with α = 0.05, power only reaches ≈ 0.3 and the false negative rate β is very high at 0.7. So it is very possible that the sizes *are* significantly different, especially since the GC 606 statistics are stable (due to the very large sample) and its coefficient of variation is stable to less than 25% of a decimated sample set. Similar to GC 606, there is signal flow from GC 9787 via coupled OFF γ+ amacrine cells to both GC 9787 itself and other short non-alpha GC dendrites in the OFF layer. Some non-alpha dendrites are themselves γ+ implying they are coupled to different sets of amacrine cells, as is the case with GC 606. While a complete tabulation of the connectivity of coupled OFF amacrine cells is over a year off in annotation time, Tulip queries reveal that some partners such as wf γ+ AC 7859 appear to be biased towards feedforward contacts, similar to IAC 9769 in the ON system.

## Discussion

### GABA signatures

GABA content is an essential signature of coupling in the ganglion cell layer. Uncoupled cells have no GABA signal but all ganglion cells have identical and inseparable glutamate signatures (Marc et al., 1990;Marc et al., 1995;Marc and Jones, 2002), regardless of GABA content (Fig.1E). As an aside, we have found no gap junctions made by any γ–ganglion cells in RC1. Ganglion cells display GABA levels ranging from undetectable to close to *bona fide* amacrine cell levels (Marc and Jones, 2002), with the majority centered around 300-600 µM, 10-fold lower than typical starburst amacrine cell levels (Fig. 1F). Given that specific ganglion cells show extensive heterocellular coupling with markers such as Neurobiotin (MW 322), it is not surprising that a molecule several times smaller such as GABA (MW 103) is highly mobile through gap junctions, similar to quantitative measures of glycine coupling into ON cone bipolar cells from glycinergic AII amacrine cells (Marc et al., 2014a). Importantly, ganglion cells known to show heterocellular coupling such as rabbit OFF alpha ganglion cells (Xin and Bloomfield, 1997a) and tON DS ganglion cells (Hoshi et al., 2009) uniformly show intermediate GABA levels (Marc and Jones, 2002). Importantly, Ackert et al., (2006;2009) and Hoshi et al., (2009) also demonstrated that an axonal cell (axon-bearing “amacrine” cell) virtually identical in dendritic morphology to our IAC 9769 was both coupled to tON DS cells and γ+ by immunocytochemistry, and that other amacrine cells with differing arbor patterns were also coupled. In contrast, ganglion cells that we know are definitively not dye-coupled such as primate midget ganglion cells (Dacey and Brace, 1992) never show GABA coupling (Kalloniatis et al., 1996).

This sets the framework for using glutamate and GABA as markers of ganglion cell coupling in other species (Fig. 2) since antibodies targeting small molecules have no species bias. In surveying our library of all vertebrate classes and many vertebrate orders (Table S2), we find that apparent heterocellular coupling between ganglion cells and amacrine cells is widespread with only one group failing to show evidence of coupling: *Trachemys scripta elegans* (formerly Genus Pseudemys), Order Testudines, Class Reptilia. As all vertebrate classes show evidence of heterocellular GC::AC coupling, this argues for such signaling as a feature of primitive retina and perhaps even of its predecessor diencephalic primordia. Indeed, extensive coupling and regions of high cell firing synchronicity is a hallmark of early mammalian brain differentiation (Niculescu and Lohmann, 2014). In sum, these considerations argue for heterocellular GC::AC coupling as a retinal *plesiomorphy*, not a synapomorphy among select classes, and that heterocellular coupling is foundational for the retina as argued by Völgyi et al., (2013a).

### Coupling and feedforward

But what is the functional network role of heterocellular coupling? A fundamental clue arose when certain retinal ganglion cells and downstream neurons in brain were found to show synchronized spiking across cells (Alonso et al., 1996;Hu and Bloomfield, 2003) and that the mostly narrowly correlated retinal firing persisted after global pharmacologic synaptic suppression, leading to the argument that it was mediated by coupling (Brivanlou et al., 1998). Subsequent analyses have refined these concepts to show that correlated spiking appears to occur within sets of the *same* ganglion cell class and that the essential pathway is heterocellular coupling with (Völgyi et al., 2013b;Roy et al., 2017). Our findings are consistent with this view, that homocellular ganglion cell coupling is either non-existent or, at best, minimal in comparison to the thousands of gap junctions each coupled ganglion cell forms with a specific array of neighboring inhibitory cells. No evidence emerged to show that any amacrine cells in the coupling networks for tON DS ganglion cells and OFF alpha ganglion cells are shared: they seem completely separate. And even though tracer studies have failed to find significant coupling among amacrine cells themselves in a given GC::AC network, we do find sparse instances of coupling among the γ+ ACs coupled to GC 606. It is somewhat remarkable that IACs are well-tracer coupled (Wright and Vaney, 2004) but show only sparse gap junctions with other candidate ACs. We have no explanation for the apparent lack of consonance between gap junction numbers and Neurobiotin visualization using photochromic 2-stage intensification as described in Vaney (1992). Perhaps IAC::IAC coupling is on more distal dendrites outside our volume.

Indeed there is a major caveat to arising from the conflicting demands of connectomics coverage and resolution. Once captured, one can downsample but never upsample, one can mine an area but never expand. Dedicating more bits to one mode steals from the other. Unlike small-field bipolar cells and glycinergic amacrine cells, ganglion cells and GABAergic neurons can have arbors much larger than a connectome. Ackert et al.,(2009) showed the axonal cells coupled to ON DS ganglion cells had fine extensions of their terminal dendrites that ascended into the OFF layer and co-stratified with the OFF ChAT+ starburst amacrine cells. As IAC 9769 has long straight dendrites that exit the full volume perimeter, such unusual morphologic features cannot be excluded. Thus there appear at least two crossover paths between the IAC and the OFF layer. In addition to the apparent cholinergic layer ramification, it is clear (via Tulip queries) that crossover glycinergic amacrine cells driven by CBa OFF bipolar cells also synapse on IAC 9769. The data of Ackert et al., (2009) do not reveal a glycinergic path, though it clearly exists. Their data support the OFF starburst amacrine cell path: OFF CBa > OFF SAC > IAC::ON DS GC; > denotes sign-conserving synapses and:: denotes coupling. It is remarkable that, despite abundant opportunity (Fig. 4), the ON γ+ IAC completely avoids the ON starburst amacrine cell arbors in favor of the OFF arbor.

IACs and ON γ+ amacrine cells coupled to tON DS ganglion cells share the same profile of excitation: a bias for class CBb4w ON cone bipolar cells and against classes CBb5 and CBb6 cells that drive starburst amacrine cells (Fig. 7). While our analysis of OFF cone bipolar cell populations is not yet as refined as for ON cone bipolar cells, the excitatory drive to amacrine cells coupled to OFF alpha ganglion cells shares similar of biases: towards OFF Cba2 bipolar cells and away from CBa1 bipolar cells. Considering the high diversity of vertebrate amacrine cell classes (Wagner and Wagner, 1988;MacNeil et al., 1999), every instance of GC::AC coupling could easily involve unique sets of amacrine cells for each coupled ganglion cell class, though such class separation may not be essential. But amacrine cells are not simply conduits for coupling. Every amacrine cell class that we know well is either GABAergic or glycinergic. Indeed, every amacrine or axonal cell involved with heterocellular GC::AC coupling whose signature can be retrieved is GABAergic. And connectomics can resolve the targets of coupled amacrine and axonal cells. Importantly, both ON and OFF instances of GC::AC coupling demonstrate feedforward directly from coupled AC to *different* classes of ganglion cells: cross-class inhibition. As schematized in **Figure 18**, heterocellular coupling allows an active ganglion cell to directly inhibit its neighbors: GC1::ON AC >_i_ GC2; >_i_ denotes sign-inverting signaling,:: denotes coupling and classes GC1 & GC2 are disjoint).

**Figure 18:**
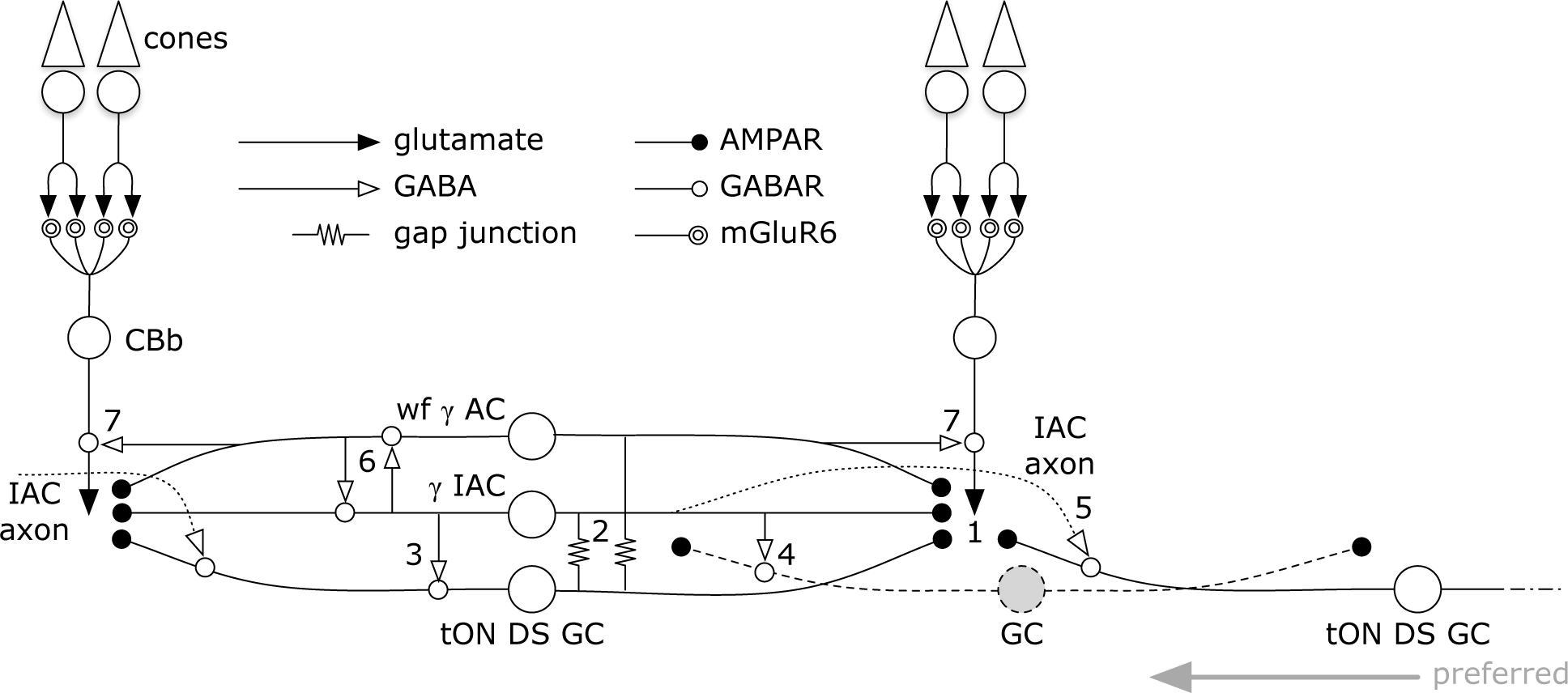
Signal flow through the tON DS GC:: γ+ AC network. Key in inset. (1) ON cone bipolar cell signals are collected by all cell classes at AMPARs or AMPARs + NMDARs. GC::AC gap junctions connect networks of (2) γ IACs and wide-field (wf) γ ACs. IACs are predominantly feedforward, driving sets of ganglion cells including (3) the coupled tON DS GC, (4) local dendrites from GCs outside this coupled set, and (5) distant instances of tON DS GCs in the far surround via their axons. IACs also engage in (6) nested feedback with wf γ ACs, which are themselves mixed feedforward (not shown) and (7) feedback inhibitory neurons. This model may also support a directional bias for tON DS GCs with the preferred direction arising from the regions driven by the axonal field of distant IACs. **Return**.

The essential feature is that inhibitory postsynaptic currents (IPSCs) should be generated in a halo of different ganglion cell classes closely synchronized with the spikes of a source ganglion cell. If these IPSCs were strong enough to suppress some incidentally coincident spikes in target ganglion cells, this could create an improved signal-to-noise ratio (SNR) at the CNS downstream targets of the source ganglion cell compared to a parallel channel (Fig. 16). Certain ganglion cells (e.g. ON-OFF DS ganglion cells) show Na-dependent dendritic spiking (Oesch et al., 2005;Schachter et al., 2010), potent spike veto by in-hibitory processes (Sivyer and Williams, 2013) and postsynaptic current integration (Brombas et al., 2017). This argues that dendritic inhibition in ganglion cells can be strong enough to suppress dendritic spiking. The bleed-through of excitation from the tON DS ganglion cell into a set of GABAergic neurons that target different ganglion cells means that such heterocellular coupling likely has the ability to suppress activity in nearby disjoint populations.

**Figure 16:**
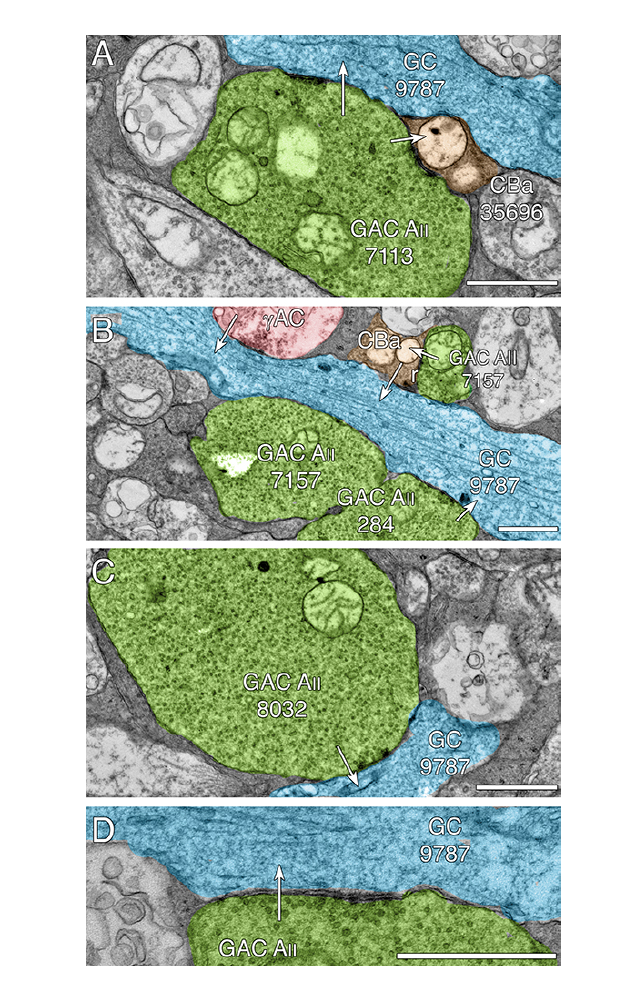
GC 9787 dendrites (cyan) collect multiple synaptic inputs from glycinergic A_II_ amacrine cell distal lobular appendages (green) across the volume. (A) Conventional synapses from GAC A_II_ 7113 onto both GC 9787 and CBa 35696 which is presynaptic to GC 9787 at two other sites. (B) Convergent signaling from γ+ amacrine cells (pink γ+ AC), GAC A_II_ 284, and a CBa bipolar cell (tan) onto GC 9787. GAC A_II_ 7157 makes synapses onto 9787 in another section (not shown) but is also presynaptic to the CBa bipolar cell. C. Single synapse from lobule GAC A_II_ 8032 onto GC 9787. D. Classical multiple presynaptic densities associated with a single GAC A_II_ synapse. Scales 1000 nm. **Return**.

While the potential for precise timing of both synchronized spikes and feedforward inhibition is clear, it is also certain that many wf γ+ amacrine cells (e.g. Fig. 12) provide feedback to cone bipolar cells of matched polarity: ON AC >_i_ ON CBb and OFF AC >_i_ OFF CBa. This provides a much broader fanout of targets for the GC::AC inhibitory couple, am-plified explicitly by the positive gain of bipolar cell ribbon synapses (e.g. Lauritzen et al., 2016).

Heterocellular coupling between spiking projection neurons and local inhibitory neurons may be more widespread than appreciated. Like retina, olfactory bulb generates synchronized oscillatory excitation / inhibition interactions that are enhanced by Cx36mediated coupling (Pouille et al., 2017) though the initial mechanism was modeled as homocellular coupling of mitral cell (MC) pools. However, detailed analysis of inhibitory intraglomerular networks provide strong evidence for heterocellular coupling between mitral cells and specific short axon GABAergic cells (SACs) in olfactory glomeruli and that the coupling, at least, is part of the mechanism that truncates events to permit more precise excitation/inhibition coordination in mitral cells (Liu et al., 2016). This may be a common mechanism in many “transient” neurons as it is consistent with feedforward onto the coupled source in both instances of ganglion cells, albeit with completely different inhibitory networks. Similarly, physiological evidence supports an analogous network for timing control in olfactory bulb. While little is known of the cell class distinctions among neighboring mitral cells in olfactory bulb, there is strong evidence for multiple projection classes, intra-bulb short-range excitatory projections, and inhibitory classes including different classes of GABAergic short axon cells (Nagayama et al., 2014). We would predict that specific classes of short axon neurons make heterocellular gap junctions with specific mitral cells and inhibit neighboring mitral cells where MC1 and MC2 are disjoint: M1::SAC>_i_ M2.

Finally, coupling networks involving inhibitory neurons can take on very complex frequency dependent operations, such as Golgi neurons in cerebellum (Vervaeke et al., 2010;Vervaeke et al., 2012;Pereda et al., 2013). In a similar fashion it is plausible that the IAC might not have effective dendritic spiking and are more passive cables, like cerebellar Golgi interneurons, but the high density of GC::AC coupling acts as an excitation repeater. Further such networks could either enhance synchrony or desynchronize in different excitation modes (Vervaeke et al., 2010).

### Direction of motion

The coupled tON DS ganglion cell is largely separate stream of directional signaling with little apparent engagement with the ON-OFF DS cohort (Ackert et al., 2006;Hoshi et al., 2011), but does contribute to the classic three-lobed orientation distribution of ON DS ganglion cells reported by Oyster and Barlow (1967). Consistent with this model we find not only complete synaptic separation of tON DS ganglion cells and the starburst amacrine cell network, but also nearly total avoidance of each other’s bipolar cell input profile. Nevertheless, like other DS cells, directional signaling is dependent on GABAergic inhibition and is suppressed after global GABA blockade (Ackert et al., 2006;Ackert et al., 2009). Directionally selectivity in visual cortex has been thought to be driven by asymmetries in excitation, although differing spatial distributions of excitation and inhibition clearly play a role (Li et al., 2017). IACs, as axonal cells, offer a built-in simple asymmetry that could be maximized for low velocity directional motion if: (1) their axons behave as classical axons and arborize into predominantly presynaptic terminals; (2) the axons do not form gap junctions; and (3) the axons target tON DS cells. We simply don’t have information on the latter in connectome RC1, but it important to consider two quantitative points. First, a complete tON DS ganglion cell likely receives over 4,000 inhibitory synapses and the bulk of those will be GABAergic (15-fold more prevalent than glycinergic synapses on GC 606). The fact that coupled ganglion cells make up a small fraction of that inhibitory input via feedforward simply means that the bulk of inhibition is driven by sets of wf amacrine cells or IAC axons arising from outside the connectome volume, displaced from the centroid of the GC 606 arbor. Importantly, prior work had shown these IACs (also known as axon bearing amacrine cells) had long sparse axons (Ackert et al., 2006;Ackert et al., 2009;Hoshi et al., 2011) but some of these are incomplete, as a more complex terminal arbor was demonstrated in Massey (2008). A more complete description of bona fide IACs in primate by Dacey (1989) describe each IAC as being surrounded by a sparse halo of axon terminals roughly 10x the diameter of the dendritic arbor and yielding perhaps over 100 fold greater coverage. Similarly, the IAC is identical to the PA1 polyaxonal cell in rabbit meticulously described by both Famiglietti (1992) and Wright and Vaney (2004). In any case, there is ample additional space in the tabulation of synapses to accommodate sparse inhibitory cells with very high coverage factors, in addition to IAC axons. Wright and Vaney (2004) also show that the net density (length) of axonal processes is ≈ 10x higher than dendritic density. If this translates to synaptic density and the axon has the same target preferences as the IAC dendrites, it is very likely that a large fraction of inhibition targeting tON DS GCs could arise from the IAC or PA1 polyaxonal cell. We have not shown that the non-IAC processes coupled to GC 606 correspond to the diffuse multistratified cell previously described (e.g. Hoshi et al., 2011), but presuming we are selecting for parts of their arbors, this cell may be even better suited than the IAC for medi-ating cross-class inhibition. The second point is a GC 606 gap junction is never more than a micron from a bipolar cell ribbon, so the shunting path for any dendritic spikes is very short. This lays the framework, via IAC axons or wf amacrine cells, for provide narrowly shaped and time-locked, delayed feedforward IPSCs to other tON DS ganglion cell instances in the coupled neighborhood.

## Conclusion

Physiological and tracer studies have firmly established heterocellular coupling as a norm in the mammalian retina. By combining small molecule markers and connectomics we provide some additional insights. First, heterocellular GC::AC coupling is likely a pleisiomorphy (a basal feature of ancient retinas) and not a synapomorphy (specialization of a clade). Second, in the instances of GC::AC coupling we know well in the mammalian retina, one involving transient ON directionally selective ganglion cells and the other engaging transient OFF alpha ganglion cells, the coupled GABAergic amacrine and axonal cells clearly inhibit many neighboring cells, including feedforward inhibition onto neighboring ganglion cells of different classes, outside the coupled set. Thus an activated ganglion cell may inhibit neighboring ganglion cells of different classes in a time-locked fashion, potentially erasing coincident dendritic spikes across ganglion cell classes. If we can now begin to tabulate and explore the detailed distributions of inhibition relative to the sites of coupling, we may uncover spatial asymmetries that convert to temporal delays necessary for encoding direction in this unique cohort of ganglion cells.

